# Allele-specific chromatin modifications of HPV-associated extrachromosomal DNAs in cervical cancer

**DOI:** 10.64898/2026.07.15.733834

**Authors:** Signe MacLennan, Michelle Ng, Vanessa L. Porter, Richard D. Corbett, Pawan Pandoh, Diane Trinh-Hamilton, Robin Coope, Yongjun Zhao, Marco A. Marra

**Affiliations:** Department of Medical Genetics, University of British Columbia, Vancouver, V6T 1Z3, Canada; Michael Smith Laboratories, University of British Columbia, Vancouver, V6T 1Z4, Canada; Canada’s Michael Smith Genome Sciences Centre, Vancouver, V5Z 1B3, Canada; BC Cancer Research Institute, Vancouver, V5Z 0B4, Canada

**Keywords:** HPV integration, extrachromosomal DNA, cervical cancer, long-read sequencing, allelic methylation, epigenetics, cancer genomics

## Abstract

Cervical cancer, a human papillomavirus (HPV) driven malignancy, is the fourth deadliest cancer in females. Approximately 25% of primary cervical cancers contain circular structures called extrachromosomal DNAs (ecDNAs), which can be entirely composed of human DNA (human-only ecDNAs) or a combination of HPV and human DNA sequences (HPV-human hybrid ecDNAs). Human-only ecDNAs harbour cancer-driving oncogenes, but epigenetic differences between human-only and HPV-human hybrid ecDNAs are unexplored. Here we report that 30% of cervical cancers contained at least one ecDNA. HPV-human hybrid ecDNAs contained more active enhancers and promoters than human-only ecDNAs and invariably contained the HPV oncogenes *E6* and *E7*. These ecDNAs also contained an allelic hypomethylation region associated with putative human promoters, as well as enrichments for motifs associated with recombination hotspots. By leveraging long-read WGS phasing data to separate ChIP-seq data into alleles, we revealed allele-specific peaks in enhancers and promoters on ecDNAs, of which HPV-human hybrid ecDNAs contained a higher density. Overall, we found HPV-human hybrid and human-only ecDNAs differed in their regulation, often at an allele-specific level. Our study more firmly establishes the importance of epigenome disruption in cervical cancer and how structural variation, particularly surrounding HPV integration sites, may shape the surrounding regulatory architecture.

## Introduction

Persistent infection with high-risk human papillomavirus (HPV; e.g. HPV16, HPV18, HPV45) drives cervical cancer^1^. Despite HPV vaccines and screening programs, cervical cancer has the fourth highest global mortality rate of cancers affecting females and is the deadliest cancer affecting females in sub-Saharan Africa^1^.

Evidence suggests the epigenome can be disrupted in cervical cancer via multiple distinct mechanisms^2,3^. For instance, cervical cancer cases are often characterized by DNA alterations in genes encoding epigenetic regulators (e.g. *KMT2D*, *CREBBP*, *EP300*)^2,4,5^. In addition, HPV integration, wherein parts of the HPV genome are incorporated into the human genome, is a frequent event in cervical cancer pathophysiology^6^, and our group and others have found that it is associated with structural variation (SV)^7,8^ and dysregulation of the surrounding epigenome^3^.

One particular SV type found in ∼25% of primary cervical cancer samples is that of extrachromosomal DNA (ecDNA)^9^. Acknowledged as a driver of cancer development and progression, ecDNA is a form of circular DNA, ranging from ∼100 kilobases (kb) to ∼5 megabases (Mb) in size, which is present at high copy number (CN) in ∼14% of primary cancer samples^9,10^. ecDNAs can serve as vehicles for oncogenes and/or their associated enhancers, enabling ecDNAs to potently drive increased oncogene expression^10–12^. Because ecDNAs appear to segregate randomly to daughter cells during mitosis^13^, they can generate intratumoural heterogeneity, which has been linked to treatment resistance^12,14^. In both pan-cancer^9^ and specific cancer contexts^15–17^, ecDNA presence has been associated with poor prognosis compared to samples lacking ecDNAs.

HPV integration can yield circular HPV-human hybrid ecDNA, which contains both HPV and human DNA^23^. Studies in cervical cancer cell lines support HPV-human hybrid ecDNAs acting as super-enhancer carriers^23^, but whether this is true in patient samples has not been investigated. There is evidence that ecDNAs containing only human DNA (i.e. human-only ecDNAs) bearing enhancers can act both in *cis*^19^ with their embedded genes and in *trans*^20^ with chromosomal genes to promote increased gene expression. However, HPV-human hybrid ecDNA chromatin regulation is unexplored. Thus, how HPV-human hybrid ecDNAs are distinguished from human-only ecDNAs in terms of their-omic features remains to be elucidated.

Here, we compared the genomic, epigenomic, and transcriptomic features of HPV-human hybrid ecDNAs to human-only ecDNAs. In general, we found that ecDNAs in cervical cancer patient samples are prevalent, have heterogeneous genomic structures, and harbour highly expressed oncogenes. We previously found regions of allelic hypomethylation on HPV-human hybrid ecDNAs^3^. Here, we report that these allelic hypomethylation regions are associated with candidate human promoters and HPV-human hybrid fusion transcript expression. We mapped allele-specific ChIP-seq data by integrating them with phasing information from long-read WGS data to show that ecDNAs often contain allele-specific ChIP-seq peaks in enhancer and promoter regions. Overall, our study finds that ecDNA structure and regulation varies depending on the presence of HPV DNA, which indicates that HPV-human hybrid and human-only ecDNAs may have different oncogenic functions.

## Results

### ecDNAs are prevalent in cervical cancer

We identified and characterized ecDNAs in a cervical cancer cohort from the Human Immunodeficiency Virus positive (HIV+) Tumour Molecular Characterization Project (HTMCP^2^; n = 118 patients, 61% HIV+) using previously generated short-read whole-genome sequencing (WGS; n = 118 patients/samples), RNA-sequencing (RNA-seq; n = 118), native chromatin immunoprecipitation sequencing (ChIP-seq; n = 50)^2^, and long-read WGS (n = 72)^3^, including previously unpublished long-read cDNA sequencing (cDNA-seq; n = 29) data (Fig. 1a). We detected candidate ecDNAs using short-read WGS data and the AmpliconArchitect (AA)^24^ software, which predicts ecDNA structures using CN changes and discordant read pair information. Using these data, we detected 50 ecDNAs in 30% (35/118) of samples (between one and four ecDNAs/sample). Of these, 37 (74%) contained only human DNA and 13 (26%) contained both HPV and human DNA (Fig. 1b).

**Figure 1.**
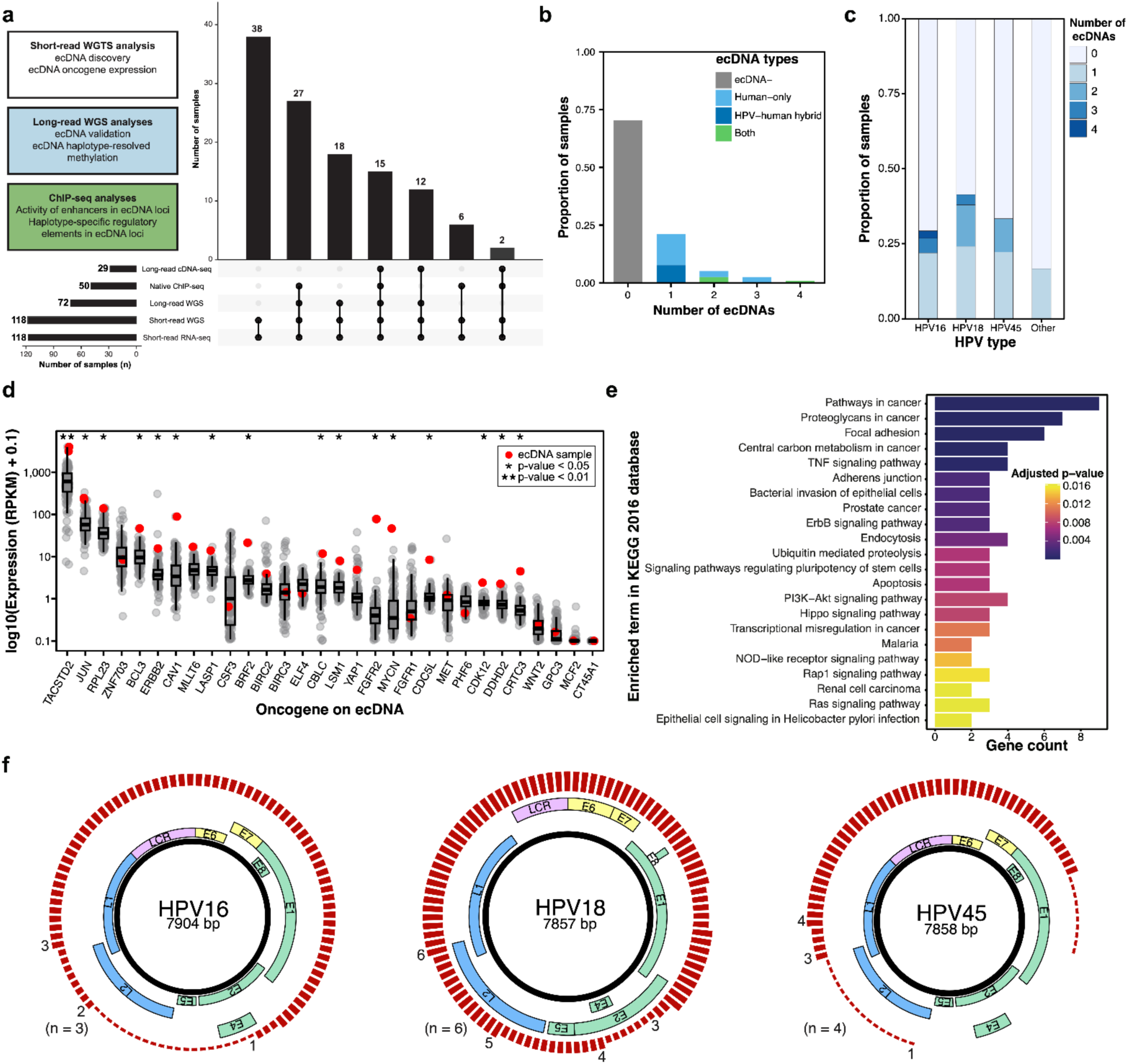
ecDNAs are prevalent in cervical cancer and often harbour highly expressed oncogenes and/or parts of the HPV genome. **a** Study design overview. **b** Proportion of cases with ecDNAs by number of ecDNAs, which refers to ecDNAs derived from distinct genomic regions within a given sample. **c** Proportion of cases with ecDNAs by HPV type. **d** Expression of oncogenes on ecDNAs and their expression relative to the rest of the cervical cancer cohort (ecDNA+ sample(s) in red n = 1-2; remaining cohort = 116-117 samples). ANOVA permutation test with false discovery rate (FDR) < 0.05 multiple testing correction, * = adj. p-value < 0.05, ** = adj. p-value < 0.01. **e** Gene set enrichment analysis (GSEA) results for oncogenes on ecDNAs. **f** Genomic regions of HPV represented on HPV-human hybrid ecDNAs where height of the red bar indicates number of ecDNAs (n = 13; with numbers over red bars indicating number of ecDNAs).

As long-read WGS is superior to short-read WGS for detecting complex SV^25^, a category which includes ecDNA, we used long read data to validate our AA results. We considered 34 candidate ecDNAs from 25 samples where long-read WGS data were available (Methods; Supp. Fig. 1a). Of the 34 candidate ecDNAs predicted using AA and short-read WGS data, we detected evidence of circularity in 29 (85%) using long-read WGS (Methods; Supp. Fig. 1b). Of these, 19 (56%) ecDNAs were assembled into circular contigs using the *de novo* long read assembler Flye^26^. Another ten ecDNAs had DNA sequence read support for circularity, as determined by viewing supplemental reads in the integrated genomics viewer (IGV)^27^ that connected contig ends to form circles (see Methods, examples in Supp. Fig. 1c-f). Five candidate ecDNAs could not be validated using either approach. These were all complex human-only ecDNAs (i.e. composed of more than one genomic segment). We excluded one human-only ecDNA from all following analyses, due to this ecDNA exhibiting multiple inconsistent structures and lacking consistent genomic coordinates. Given that our long-read WGS validation results were generally consistent with AA results, we concluded that AA-predicted ecDNAs were generally reasonably well estimated across our samples (n = 118).

### General ecDNA structural features and correlates

We observed single ecDNAs in 25/35 (71%) of ecDNA+ cases. In 10/35 (29%) ecDNA+ samples, we observed between two and four distinct ecDNAs derived from different genomic regions (Fig. 1b). Interestingly, although ten samples contained multiple ecDNAs, we never observed more than one HPV-human hybrid ecDNA within a sample. The proportion of samples that were ecDNA+ varied slightly depending on the HPV type of the sample, ranging from 17% in the case of rare HPV types (i.e. HPV types other than HPV16, HPV18, or HPV45) to 41% for HPV18 (Fig. 1c).

Next, we looked for molecular correlates of ecDNA presence across the cohort. We first considered significantly mutated genes (SMGs), which have higher mutation rates than the background rate and are therefore candidate cancer drivers^28^. Analysis of SMGs across the cohort using dNdScv^29^ did not reveal obvious correlations of drivers with ecDNA+ samples (Supp. Fig. 2a, see Methods). We also examined the relationship between molecular and clinical covariates (i.e. HIV status, clinical stage, histology, APOBEC signature, HRD, and HPV clade) and ecDNA classes (i.e. human-only, HPV-human hybrid, or both) and none of these covariates correlated with the ecDNA classes across samples (Supp. Fig. 2a).

We next sought to relate ecDNA classes to mutational signatures predicted by MuSiCal^30^ (using SNVs called by Strelka2^31^), as human-only ecDNAs in other cancer contexts have been linked to APOBEC signatures^32^. *De novo* signature discovery uncovered four signatures, which matched the COSMIC signatures associated with defective homologous recombination DNA damage repair (SBS3), ultraviolet light exposure (SBS7a), activity of APOBEC family of cytidine deaminases (SBS13), and unknown (SBS39; Supp. Fig. 2b). Notably, the proportion of samples containing evidence of each signature varied based on ecDNA class (Supp. Fig. 2c), with HPV-human hybrid ecDNA-containing samples having the highest proportion of samples affected by SBS3, SBS13, and SBS7a.

As ecDNAs are vehicles for oncogenes^19^, we next investigated the genic content of ecDNAs in our cohort. In total, AA predicted 444 unique human genes residing on ecDNAs, including 33 different oncogenes. Of these oncogenes, 30 genes had apparently complete coding sequences, and five genes (*BCL3*, *CBLC*, *ERBB2*, *JUN*, and *TACSTD2*) were detected on ecDNAs in two separate samples (Supp. Table 1, see Methods). 16/33 (48%) oncogenes had significantly higher expression in the sample predicted to contain an ecDNA bearing that oncogene compared to other samples from the cohort (ANOVA permutation test; FDR < 0.05; Fig. 1d). We performed a gene set enrichment analysis for oncogenes on ecDNAs and uncovered a significant enrichment in metastasis-related genes (e.g. focal adhesion: adj. p-value = 1.8 x 10^-5^) and *ERBB* family members (adj. p-value = 3.3 x 10^-3^; Fig. 1e). Notably, *ERBB* family members are often found on ecDNAs in other cancers, for example *EGFR* in glioblastoma^12^.

All HPV-human hybrid ecDNAs contained the HPV oncogenes *E6* and *E7* (13/13; 100%). These ecDNAs infrequently contained an intact *E2* (3/13; 23%; Fig. 1f), which encodes a negative regulator of *E6* and *E7* and is commonly lost or disrupted in the process of HPV integration^33,34^. Notably, all HPV-human hybrid ecDNAs also contained the HPV long control region (LCR), which contains the viral origin of replication and plays a vital role in positively regulating transcription of *E6* and *E7*^35^.

### ecDNA genes tend to be mono-allelically expressed

Next, we sought to distinguish the expression of ecDNA human genes from the same genes in their normal chromosomal contexts. As ecDNAs were derived from only one allele^37^, we focused on characterizing mono-allelic vs bi-allelic expression patterns. We used the computational tool IMPALA^36^ on our short-read RNA-seq data that had matched long-read WGS data (n = 46 samples; Methods), confining our analysis to ecDNA genes that had sufficient numbers of heterozygous SNVs to unambiguously assign transcripts to alleles (“phasing”). This filter retained 95 of the 444 genes identified using AA. We confirmed that genes on ecDNAs tended to exhibit allele specific expression (ASE; Supp. Table 2), in contrast to these same genes when they were not on an ecDNA (Fisher’s exact test p-value < 2.2 x 10^-16^; Supp. Fig. 3a). For example, when a gene was on an ecDNA in a given sample, that gene tended to have ASE in that sample, whereas that same gene in ecDNA-samples was significantly less likely to have ASE in those ecDNA-samples.

To understand the mechanisms contributing to ASE of ecDNA genes, we investigated the methylation status of gene promoters. Surprisingly, promoter methylation frequency was higher in ASE genes compared to BAE genes regardless of whether the gene was predicted to reside upon an ecDNA, indicating that promoter hypomethylation does not drive ASE in this cohort (Welch’s two-sided t-test p-value = 9.5 x 10^-5^; FDR < 0.05; Supp. Fig. 3b). Notably, although less common, promoter hypermethylation is associated with gene activation in certain cancer contexts^37^. Although allele-specific promoter methylation differences were greater for ASE genes compared to BAE genes (Supp. Fig. 3c), the differences were small (∼0.01 difference in methylation frequency between alleles), leading us to consider that the ASE of ecDNA genes may be driven by other factors, e.g. their high CN, rather than methylation differences between alleles. Interestingly, annotated oncogenes on ecDNAs tended to have higher ASE compared to non-oncogenes on ecDNAs (Two-sided Mann-Whitney *U* test adj. p-value = 0.036; FDR < 0.05; Supp. Fig. 3d), but oncogenes did not have lower promoter methylation values (Two-sided Mann-Whitney *U* test adj. p-value = 0.55; FDR < 0.05; Supp. Fig. 3e). Overall, this observation is compatible with the notion that apparent non-oncogenes may be passengers on ecDNAs, and are not under the same selective pressures as oncogenes, at least in terms of expression.

### A subset of ecDNAs contain aDMRs overlapping gene promoter regions

To better understand the global regulation of ecDNAs, we investigated ecDNA-specific methylation patterns in more detail. When excluding allele information, we found no significant difference in ecDNA methylation in samples predicted to contain ecDNAs and either those same loci in ecDNA-samples (n = 49 samples total) or a random sample of similarly sized genomic regions (n = 100, see Methods; Supp. Fig. 4a-b). However, we could not exclude the possibility that ecDNA methylation is dysregulated on an allele-specific basis.

To investigate ecDNA-specific methylation patterns using allele-resolved data, we first asked whether ecDNAs contained an enrichment of allelic differentially methylated regions (aDMRs) relative to a null distribution (see Methods). Only 1/18 (5.5%) of ecDNAs with available long-read WGS data were enriched in aDMRs, leading us to conclude that ecDNAs are generally not enriched in aDMRs relative to the rest of the genome (Supp. Fig. 4c). Furthermore, we found that the majority of aDMRs overlapped repetitive regions (143/218, 66%) and that less than half of the aDMRs overlapped genes (95/218, 44%) or gene promoters (18/218, 8.3%, Supp. Fig. 5a; Supp. Table 3).

We also sought to understand which genes contained aDMRs on ecDNAs and if ecDNA types varied in predicted aDMR frequency. Consistent with there being more human-only ecDNAs in our cohort and the size of human-only ecDNAs being significantly larger than HPV-human hybrid ecDNAs, we found that the majority of ecDNA aDMRs were in human-only ecDNAs (Supp. Fig. 5b-e). A number of larger genes had multiple aDMRs falling within them on ecDNAs (Supp. Fig. 5c-d), and *BCL3*, *CDH5*, *ANKRD7*, *CTTNBP2*, *C7orf65*, and *NLGN4X* had more than one predicted promoter aDMR in an ecDNA locus (Supp. Fig. 5e). These observations indicated that ecDNA methylation is not disrupted at a global level nor in a consistent manner, but that allele-specific methylation profiles still exist in a subset of ecDNAs, of which a subset overlap gene promoter regions.

### Human oncogenes on ecDNAs are hypo-methylated on both alleles

Given that oncogene promoters have been reported to be hypomethylated in an ecDNA-specific manner^22^, we next tested whether oncogene promoters exhibited allele-specific hypomethylation in ecDNA loci. We hypothesized this would not be observed in our cohort, as we did not uncover any aDMRs falling within annotated oncogene promoter regions. We found that the ecDNA oncogene promoters showed very little difference in methylation profiles between alleles, supporting our earlier aDMR analysis results (Supp. Fig. 5f). However, we did observe that oncogenes residing in ecDNA loci tended to be hypomethylated on both alleles and that this effect was not ecDNA-sample specific (ANOVA permutation test adj. p-value = 0.0020; FDR < 0.05), perhaps indicating that ecDNA formation may selectively affect actively transcribed oncogenes (Supp. Fig. 5g). Interestingly, the *CDK12* locus was an outlier to this trend, as the promoter regions for *CDK12* were highly methylated in the ecDNA-containing sample, but also on both alleles (ANOVA permutation test adj. p-value = 0.0020; Supp. Fig. 5g).

### HPV-human hybrid ecDNAs are smaller than human-only ecDNAs, and harbour fewer genes but more structural variation

We next investigated whether HPV-human hybrid ecDNAs are structurally distinct from human-only ecDNAs by comparing the genic and repetitive element features of human-only (n = 36) and HPV-human hybrid (n = 13) ecDNAs. We found that HPV-human hybrid ecDNAs were significantly smaller than human-only ecDNAs (median size of ∼77 kb vs ∼232 kb; Two-sided Mann-Whitney *U* test p-value = 1.5 x 10^-2^; Fig. 2a) and had a tighter distribution of sizes (standard deviation [*σ*] = 80 and 2,700, respectively). HPV-human hybrid ecDNAs also contained significantly fewer human genes than human-only ecDNAs (Two-sided Mann-Whitney *U* test p-value = 2.8 x 10^-2^; Fig. 2b; see Methods). Notably, ecDNA size and the number of human genes contained per ecDNA were significantly correlated for human-only ecDNAs (R = 0.83; Pearson correlation p-value = 3.5 x 10^-10^), but not correlated among HPV-human hybrid ecDNAs (R = 0.17; Pearson correlation p-value = 0.59; Fig. 2d), perhaps indicating HPV-human hybrid ecDNAs are not serving as vehicles for human genes, although the sample size was low (n = 13).

**Figure 2.**
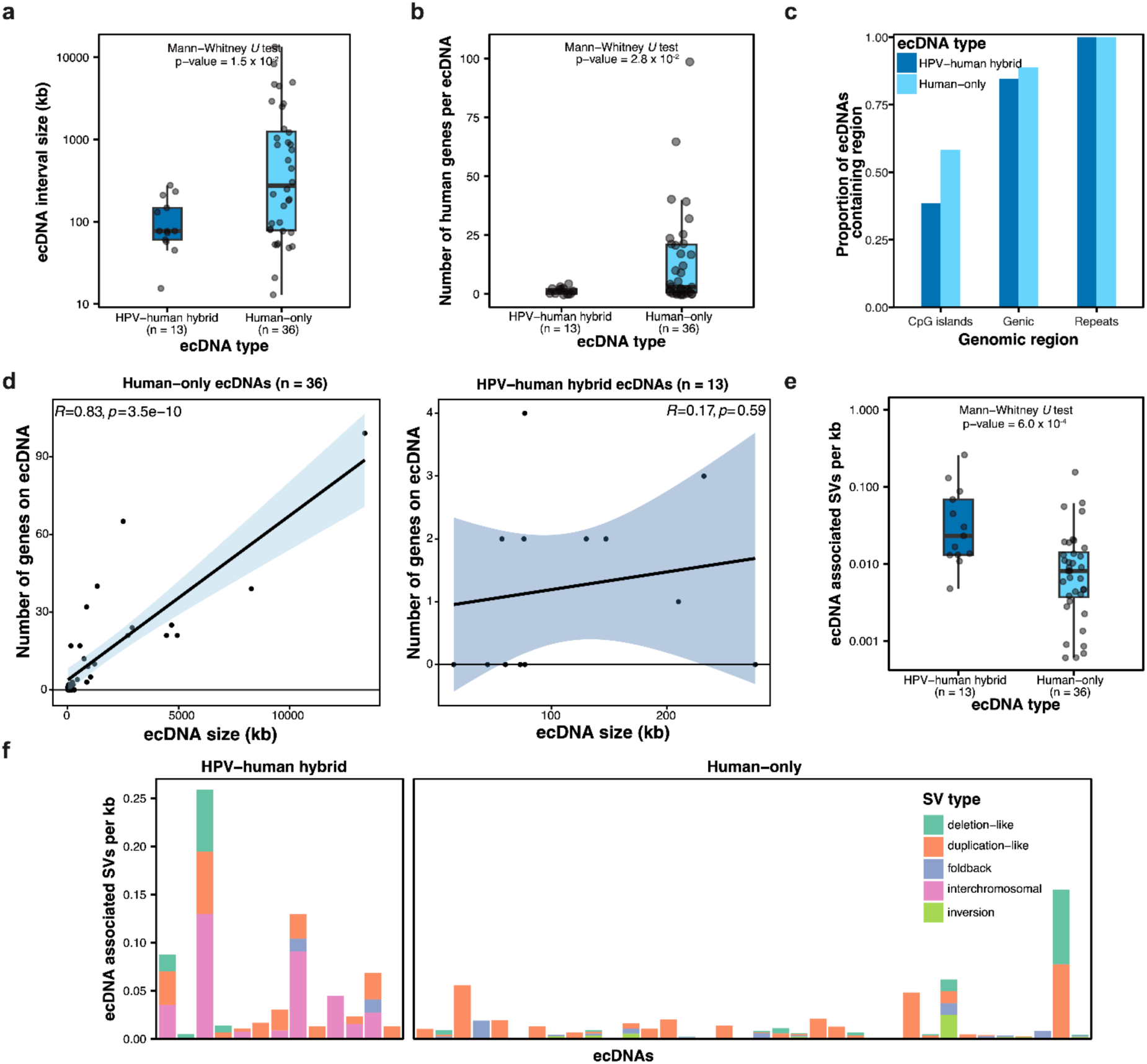
Human-only ecDNAs are larger and contain more human genes, but less structural variation than HPV-human hybrid ecDNAs. **a** ecDNA interval size comparing ecDNA types. **b** Number of human genes per ecDNA. **c** Proportion and types of genomic segments composing ecDNAs. **d** Correlation of ecDNA size (in kb) and number of human genes residing on the ecDNA. **e** ecDNA associated SVs per kb comparing ecDNA types. **f** Number and types of ecDNA-associated SVs normalized for ecDNA size (for SV definitions see Ho et al. 2020^61^ or Smolka et al. 2024^62^).

Regardless of ecDNA type, all ecDNAs contained repetitive sequences and most contained regions annotated as genic (32/36; 88.9% of human-only and 11/13; 84.6% of HPV-human hybrid ecDNAs; Fig. 2c). Neither ecDNA type was derived from chromosomes 9, 20, or 22. We next compared the frequency of different repetitive elements on ecDNAs (Supp. Fig. 6a, see Methods). When compared to a random sample of similarly sized genomic regions (n = 9,097), ecDNAs (n = 49) contained significantly more repeats per kb (Welch’s Two Sample t-test p-value = 5.0 x 10^-5^; Supp. Fig. 6b). However, after multiple testing correction no specific repeat class was enriched on ecDNAs as compared to this random sample of similarly sized genomic regions (Welch’s Two Sample t-test; FDR < 0.05; Supp. Fig. 6c). Repetitive regions on ecDNAs have not been extensively studied, but a recent report suggests repeats, in particular transposable elements on ecDNAs, may play a role in facilitating gene activation^38^.

We also compared ecDNA SVs (i.e. SVs within ecDNA regions; see Methods) and found that HPV-human hybrid ecDNAs contained a higher frequency of SVs per kb compared to human-only ecDNAs (two-sided Mann-Whitney *U* test p-value = 6.0 x 10^-4^; Fig. 2e). However, in general, human-only ecDNAs exhibited a wider diversity of SV types (Fig. 2f), which may be due to different ecDNA formation mechanisms between ecDNA types.

Interested particularly in ecDNA formation, we next investigated whether ecDNAs were associated with chromothriptic regions of the genome and whether this varied by ecDNA type. To do this, we analysed long-read WGS data (n = 72) using ShatterSeek^39^, which detects chromothripsis based on copy number oscillation and SV clustering criteria. Overall, 12/72 (17%) of samples were associated with at least one chromothripsis event (median 1, range 1-2), which is slightly lower than previously reported literature for cervical cancer, which puts that figure closer to 25%^39^. We also found that no ecDNA region overlapped a chromothripsis event and ecDNA+ samples were not significantly associated with chromothripsis presence compared to ecDNA-samples (Fisher’s exact test p-value = 0.18).

### Allelic methylation state associated with HPV integration and HPV-human hybrid ecDNAs is associated with a putative human promoter and fusion transcript expression

We generated high-quality long-read cDNA-seq data for 29 samples, of which 24 were ecDNA+ (median read count of 78 million reads, yield of 46 Gb, N50 of 650 bp, and chimerism rate of 0.80%; Supp. Fig. 7). Notably, the percentage of reads representing annotated full-length transcripts (Methods) was low (mean of 0.87%, Supp. Fig. 7f), but were long enough to identify evidence for human-HPV chimeric fusion transcripts.

By integrating long-range phasing information from the long-read WGS data (see Methods), we found examples of allelic methylation differences in human DNA segments where HPV integration and HPV-human hybrid ecDNA formation was predicted. Of the nine samples with HPV-human hybrid ecDNAs, eight had sufficient long-read WGS read coverage (one had phasing errors in the ecDNA locus) and five of these (62.5%) had evidence of allelic methylation differences downstream of the HPV integrant (example in Fig. 3a-b). Notably, all eight HPV-human hybrid ecDNAs were transcribed (Fig. 3c). In all five cases with allelic hypomethylation, the predicted HPV-human ecDNA allele was hypomethylated downstream of HPV transcription. Other samples lacking the particular HPV-human hybrid ecDNA tended to be methylated on both alleles. Notably, other forms of HPV integration were found to have inconsistent human methylation patterns immediately up-and down-stream of HPV integrants^3^. Overall, these results are compatible with at least the notion that HPV integration, resulting in HPV-human hybrid ecDNA formation, may be more likely to occur in regions of pre-existing allelic hypomethylation. Alternatively, HPV integration may induce hypomethylation during the process of HPV-human hybrid ecDNA formation.

**Figure 3.**
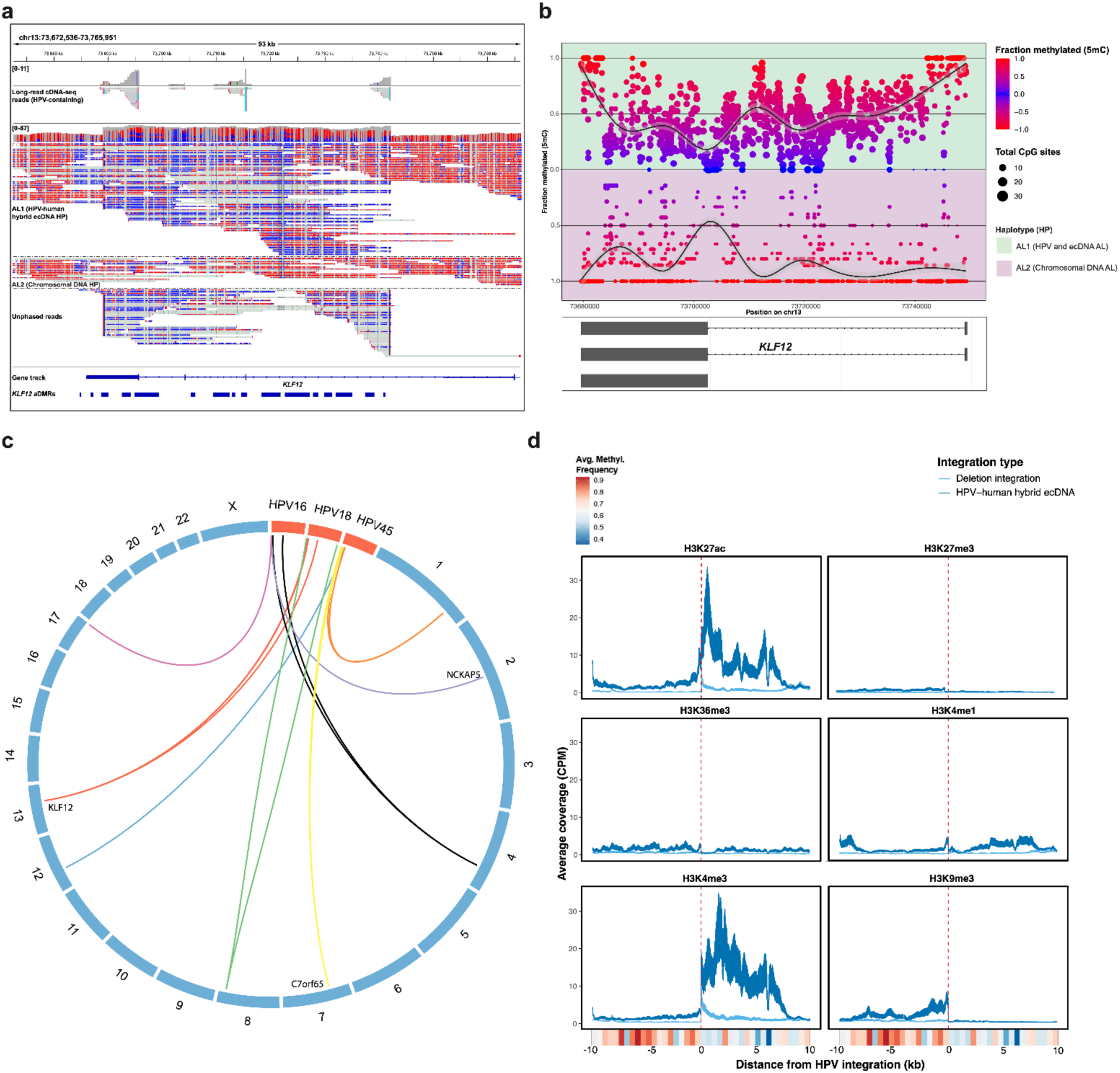
HPV-human hybrid transcripts from ecDNA loci and putative human promoters are associated with allele-specific hypomethylation. **a** Example of an IGV picture of regions of allelic hypomethylation associated with HPV integration and HPV-human hybrid ecDNA formation. **b** Methylation frequency diagram showing a zoomed in version of b. **c** HPV-human hybrid ecDNA integration breakpoints detected in long-read cDNA-seq data (each colour represents a different ecDNA/sample). **d** Average ChIP-seq coverage per basepair for 10 kb up-and down-stream of HPV integration on HPV-human hybrid ecDNAs (n = 8) compared to deletion integrations described in Porter et al. 2024^3^. Average binned methylation (bin size = 500 bp) frequencies are shown below in the coloured bars. DMR = differentially methylated region (allelic), AL = allele.

To assess the histone modification landscape surrounding HPV integrants, we also analyzed six histone mark profiles (H3K27ac, H3K4me1, H3K4me3, H3K9me3, H3K36me3, H3K27me3) from samples containing HPV-human hybrid ecDNAs. Analysis of regions 10 kb up-and down-stream of HPV integrants in these samples (n = 8), revealed the average signal across all samples was indicative of a putative DNA promoter within the human DNA immediately downstream of HPV integrants on HPV-human hybrid ecDNAs. This was also in contrast to deletion-type integrations found in other samples in this cohort (Fig. 3d; as we described previously^3^). The average predicted human promoter was also hypomethylated on the ecDNA allele, as noted in the previous analysis (Fig. 3d).

### Human genes in allelically hypomethylated regions on HPV-human hybrid ecDNAs tend to exhibit elevated and allele-specific expression

Next, we analyzed the expression patterns of human genes falling within regions of allelic hypomethylation downstream of HPV on HPV-human hybrid ecDNAs. To probe expression from HPV-human hybrid ecDNAs, we utilized our ASE and long-read cDNA-seq data.

Of the five samples with allelic hypomethylation downstream of HPV on an HPV-human hybrid ecDNA, three encompassed parts (*KLF12*, *NCKAP5*) or entire human genes (*C7orf65*). Notably, all three genes had ASE in the samples predicted to contain an allelic hypomethylation region downstream of HPV integration. *KLF12* also had its highest expression in the sample containing a HPV-human hybrid ecDNA as compared to the rest of the cervical cancer cohort, although its expression across the cervical cancer cohort was not elevated compared to the Genotype-Tissue Expression (GTEx) project^40^. *NCKAP5* expression was not elevated in the ecDNA-containing sample, but we note it is a large gene (∼1 Mb), and the allelic hypomethylation region was in an intronic region.

We next focused on *C7orf65*, as the allelic hypomethylation region encompassed the entire gene (Fig. 4a-c). Interestingly, *C7orf65* exhibited low abundance not only in the ecDNA-containing sample, but in approximately half the cohort (58/118, 49%), whereas it was generally not expressed in normal tissues (n = 19; Two-sided Mann-Whitney *U* test p-value = 8.5 x 10^-4^; Fig. 4d). Although the function of *C7orf65* is not well understood^41^, its cancer-enriched expression may indicate a role in cervical cancer pathobiology. We didn’t observe significant allele-specific methylation differences in *C7orf65* within the other cervical cancer samples in our cohort, suggesting the regulation of *C7orf65* expression via methylation may be a minor and potentially ecDNA-specific mechanism of expression regulation (Fig. 4e).

**Figure 4.**
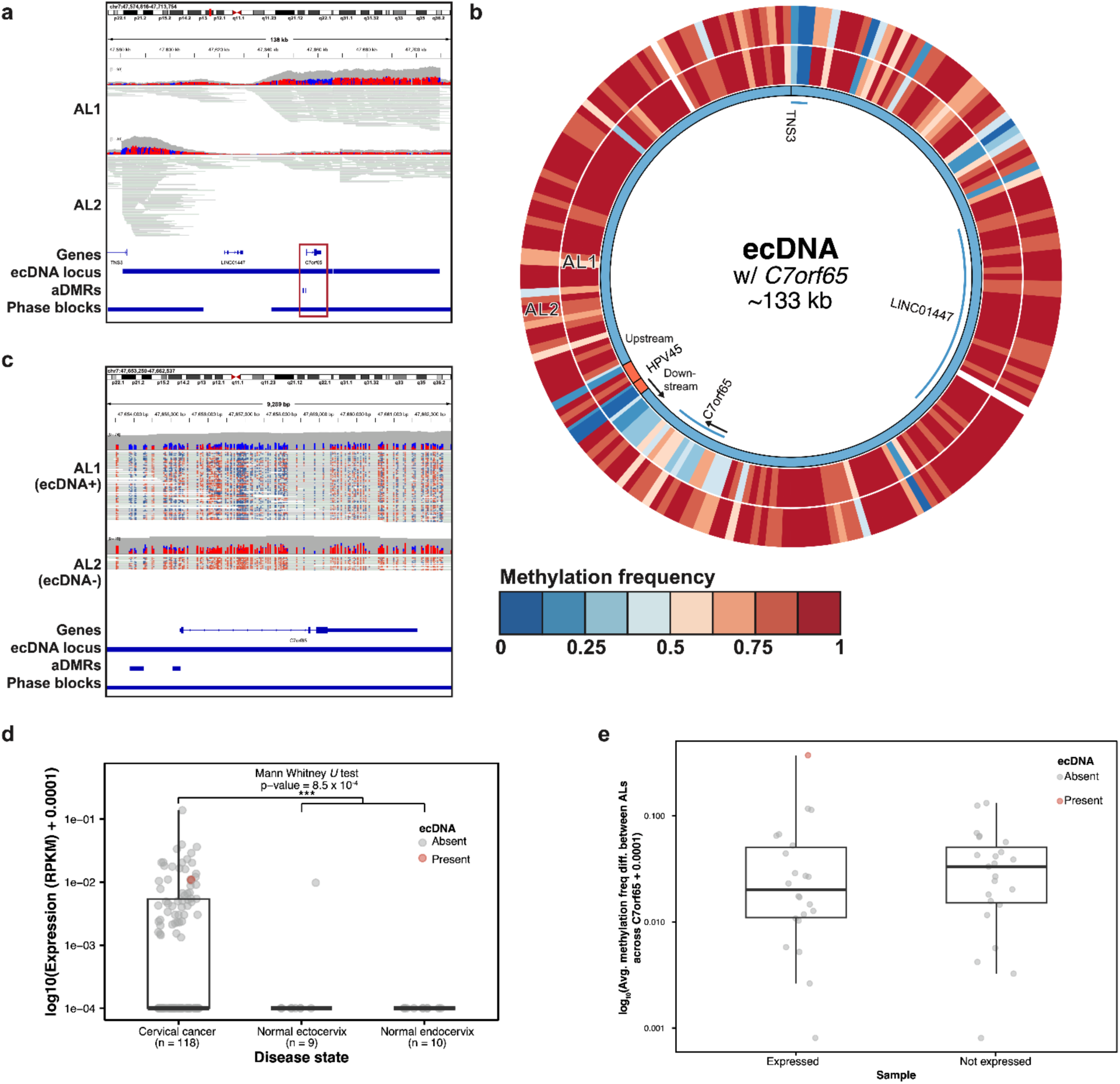
An example of allelic differentially methylated regions (aDMRs) overlapping a gene promoter (*C7orf65*) within an ecDNA locus. a All methylation phased reads covering the ecDNA locus, which encompasses the *C7orf65* gene (methylated CpGs in red, unmethylated CpGs in blue). **b** Circos plot showing predicted ecDNA structure from AmpliconArchitect and supported by long read validation and allele-specific methylation. Note that HPV45 is not allelically resolved (all HPV45 reads are included and not separated into alleles). **c** Zoomed in segment of ecDNA highlighting the aDMR covering *C7orf65* (as indicated by red box in **a**). **d** RNA expression of *C7orf65* across our cervical cancer cohort (n = 118) and compared to normal endocervix and ectocervix data from GTEx. **e** Average methylation frequency across *C7orf65* gene for all samples in our cohort stratified by *C7orf65* expression status (ANOVA permutation test with FDR < 0.05 multiple testing correction,* = adj. p-value < 0.05, ** = adj. P-value < 0.01, *** = adj. p-value < 0.001.

### HPV-human hybrid ecDNAs are enriched in human promoters and viral enhancers

Super enhancers have been reported on HPV-human hybrid ecDNAs^23^. We thus used ChIP-Seq data from 50 samples, seeking to overlay H3K27ac and H3K4me1 peaks and H3K27ac and H3K4me3 peaks to identify *de novo* enhancers and promoters respectively in predicted ecDNA regions (see Methods). In this analysis, we could not isolate ecDNA-specific ChIP-seq signals, but nonetheless could analyze peaks falling within predicted ecDNA regions.

To answer whether ecDNAs are enriched in enhancers and/or promoters and whether this varies depending on ecDNA type, we first compared human DNA regions on HPV-human hybrid ecDNAs and human-only ecDNAs to each other and to these same regions in ecDNA-samples. We observed that HPV-human hybrid ecDNAs had a higher density of human promoters when compared to both human-only ecDNAs (Welch’s two Sample t-test adj. p-value = 0.015; FDR < 0.05) and matched regions in ecDNA-samples (Welch’s two Sample t-test adj. p-value = 0.015; FDR < 0.05, Fig. 5a). There was also a non-significant trend in which HPV-human hybrid ecDNAs had a higher density of enhancers compared to human-only ecDNAs (Welch’s Two Sample t-test adj. p-value = 0.051; FDR < 0.05; Fig. 5a). When focusing on the segments of HPV DNA present on HPV-human hybrid ecDNAs, we found a significant enrichment for predicted enhancers on HPV-human hybrid ecDNAs compared to these viral DNA segments in other samples lacking these ecDNAs (Welch’s Two Sample t-test p-value = 0.024, Fig. 5b). When controlled for the size of human DNA regions on ecDNAs, HPV-human hybrid ecDNAs tended to have increased total human promoter signal compared to human-only ecDNAs (as measured by the sum of CN-adjusted coverage of H3K4me3 across the human DNA regions of each ecDNA; Welch’s two Sample t-test adj. p-value = 0.048; FDR < 0.05; Fig. 5c). Notably, HPV integration events tend to occur in predicted active and open chromatin regions, including those marked by H3K4me3 and H3K27ac^42^. Interestingly, samples containing HPV-human hybrid ecDNAs had significantly higher E6 expression and lower E2 expression (Fig. 5d), which may be due to HPV-human hybrid ecDNAs having a more permissive chromatin state compared to other forms of HPV integration.

**Figure 5.**
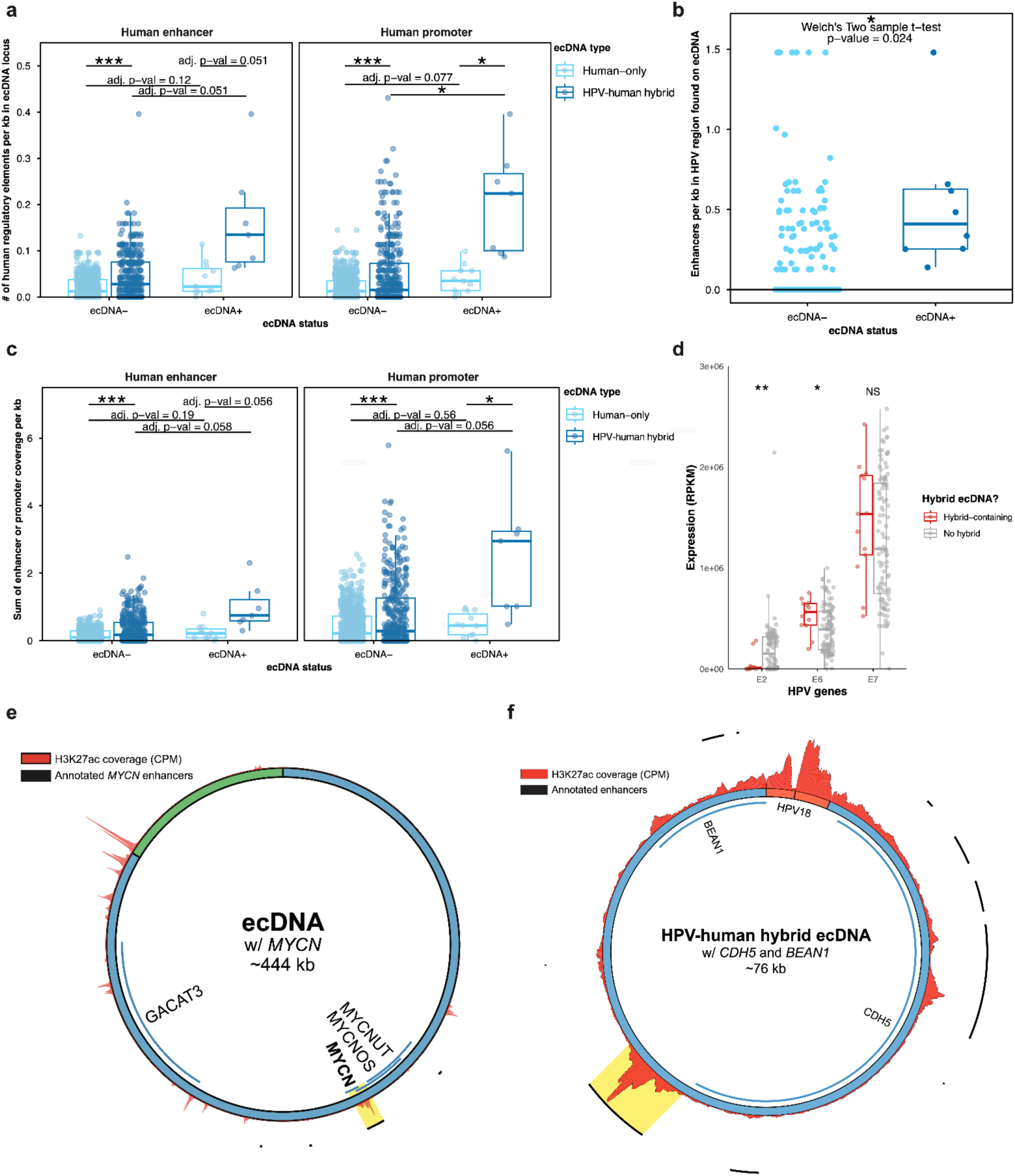
HPV-human hybrid ecDNAs contain a higher density of human promoters and viral enhancers. **a** Human enhancers and promoters per kb (*de novo* discovery), for HPV-human hybrid and human-only ecDNAs. **b** Viral enhancer density in HPV regions on HPV-human hybrid ecDNAs compared to these same regions in ecDNA-samples. **c** Sum of coverage (proxy for activity) of human enhancers and promoters in ecDNA loci. **d** RNA expression for samples containing HPV-human hybrid ecDNAs (n = 13) compared to those lacking HPV-human hybrid ecDNAs (n=105). **e** H3K27ac coverage for four annotated *MYCN* enhancers within an ecDNA locus. Enhancers are indicated by black lines surrounding the ecDNA circle. Enhancer overlapping the *MYCN* oncogene is highlighted in yellow. **f** H3K27ac coverage for ten annotated enhancers within a HPV-human hybrid ecDNA locus. Enhancers are indicated by black lines surrounding the ecDNA circle. The enhancer with the highest H3K27ac coverage is highlighted in yellow.

We next used Genehancer^43^ annotations and H3K27ac ChIP-seq signals to investigate the presence of annotated enhancers contained within ecDNA loci (see Methods). Here, in an allele agnostic manner, we define an ecDNA locus as the genomic region from which an ecDNA arose. Although H3K27ac coverage is generally increased across ecDNAs, likely as a result of high CN, H3K27ac coverage varied across ecDNAs for both human-only ecDNAs (example in Fig. 5e) and HPV-human hybrid ecDNAs (example in Fig. 5f), with apparent peaks sometimes, but not always, corresponding to annotated enhancers. ecDNAs often contain annotated enhancers that activate oncogenes also found on the ecDNA, such as *MYCN* (Fig. 5e) or genes with apparent oncogenic functions, such as *CDH5* and *BEAN1* (Fig. 5f). Notably, *MYCN* enhancers, have been reported as an example of ecDNA-mediated enhancer hijacking in neuroblastoma^11^, *CDH5* plays roles in cell-to-cell adhesion and angiogenesis^44,45^, and high *BEAN1* expression correlates with poor prognosis in rectal adenocarcinoma^46^.

Altogether, HPV-human hybrid ecDNAs were enriched in human promoters and viral enhancers suggesting that HPV integration may affect histone modification landscapes on ecDNAs.

### HPV-human hybrid ecDNAs are enriched in PRDM9 recombination hotspot motifs

Next, to gain potential insights into ecDNA formation and regulation, we investigated motifs that were enriched on HPV-human hybrid ecDNAs, identifying motifs enriched in enhancers (H3K4me1+H3K27ac) and H3K4me3+H3K36me3 marked regions on HPV-human hybrid ecDNAs (n = 8 HPV-human hybrid ecDNAs). Notably, H3K4me3 marks promoter regions, whereas H3K36me3 marks gene bodies, so these marks are often mutually exclusive^47^. However, overlaps in these two marks can be found at promoter/gene body boundaries, which can be recombination hotspots during meiosis^47^. To expand our search space, we looked at whether any motifs we previously found as enriched in our analysis of ChIP-seq peak overlaps were found more generally on other ecDNAs lacking ChIP-seq data by looking at motif enrichments for the entire predicted ecDNA regions (n = 13 HPV-human hybrid ecDNAs).

We detected one 20-bp long motif, MA1723.2, which was enriched 10-fold (Fig. 6a-b, Welch’s two-sample t-test p-value = 2.1 x 10^-3^; Methods) in both our H3K4me1+H3K27ac and H3K4me3+H3K36me3 screens in HPV-human hybrid ecDNA loci (Fig. 6a; see Methods for filtering strategy). MA1723.2 is a PRDM9 motif associated with recombination hotspots and double-stranded breaks^48,49^. Of particular relevance was its enrichment in the HPV-human hybrid ecDNA H3K4me3+H3K36me3 screen, as PRDM9 is a H3K4me3/H3K36me3 writer protein^47^, but this motif was also enriched more generally across all 13 HPV-human hybrid ecDNA loci. The relevance of a PRDM9 motif being enriched in enhancer (H3K4me1+H3K27ac marked) regions is unknown. Among H3K4me3+H3K36me3 overlapping peak regions in HPV-human hybrid ecDNAs loci, MA1723.2 was enriched ∼10 times among these regions as compared to nucleotide and size matched background regions (Welch’s two-sample t-test p-value = 2.1 x 10^-3^).

**Figure 6.**
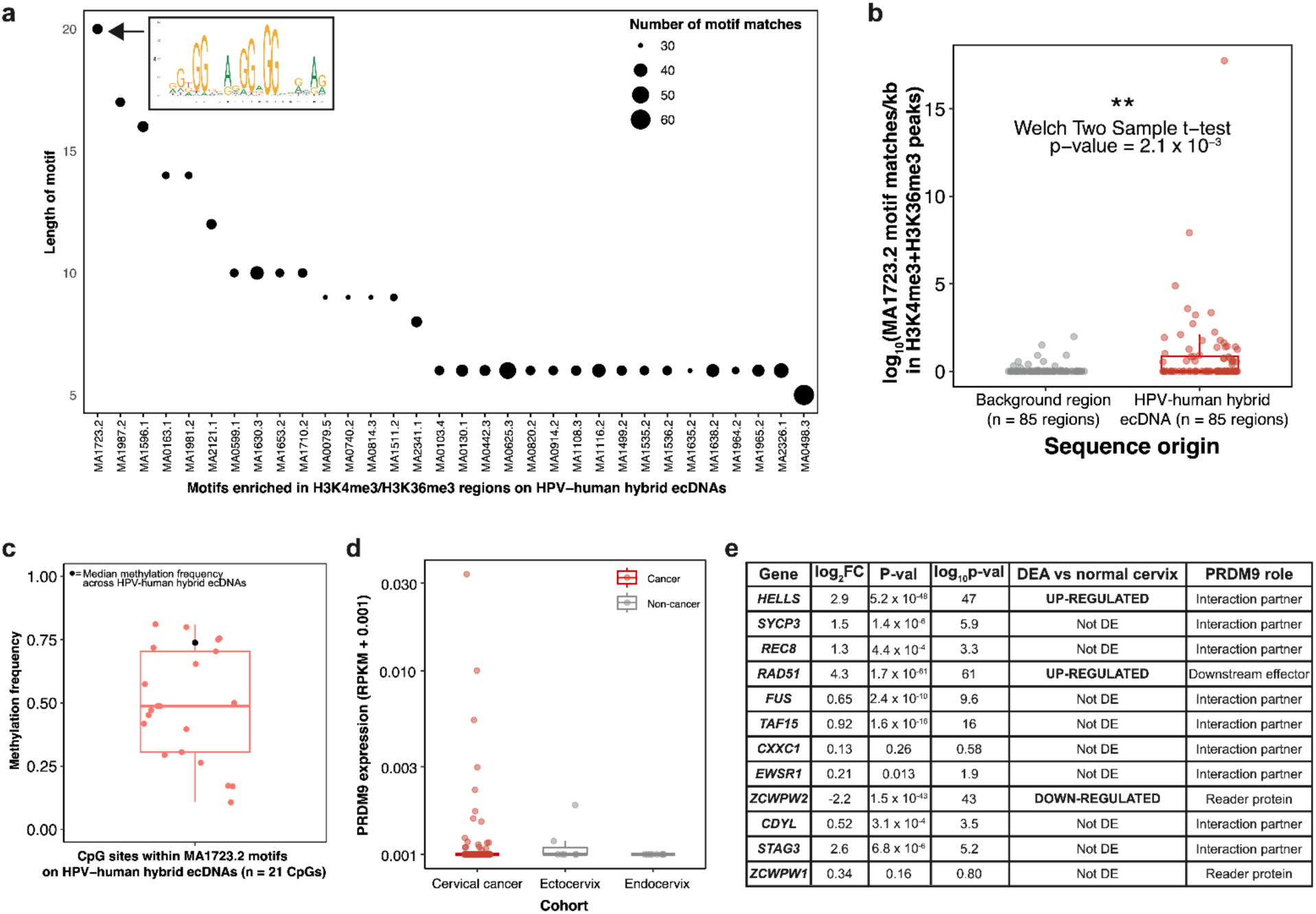
HPV-human hybrid ecDNAs are enriched in PRDM9 recombination hotspot motifs. **a** Motifs enriched at H3K4me3/H3K36me3 marked regions in HPV-human hybrid ecDNA loci with at least 30 matches among the tested regions (n = 85 regions). **b** Enrichment of MA1723.2 (PRDM9 motif) on HPV-human hybrid ecDNAs in H3K4me3/H3K36me3 regions as compared to background regions with matched nucleotide frequencies and sizes. **c** Methylation frequencies among CpG sites found within motif matches for MA1723.2 on HPV-human hybrid ecDNAs H3K4me3/H3K36me3 regions. **d** RNA expression of PRDM9 in the HTMCP cervical cancer cohort (n = 118) as compared to GTEx (n = 18). **e** Differential expression analysis (DEA) results for PRDM9 related genes.

PRDM9-like motifs on HPV-human hybrid ecDNAs within H3K4me3+H3K36me3 marked regions were slightly hypomethylated at few CpG sites (Fig. 6c; median methylation frequency of 0.49, n = 21 CpGs) as compared to HPV-human hybrid ecDNA loci as a whole (median methylation frequency of 0.74, n = 8,400 CpGs). In the short-read RNA-seq data, PRDM9 was expressed at a low level in 15% (18/118) of the cervical cancer cohort. This was not significantly different than the proportion of normal cervix samples that expressed PRDM9 (11% of normal cervix samples in GTEx, Welch’s Two Sample t-test p-value = 0.21) and did not overlap with HPV-human hybrid ecDNA presence (Fig. 6d; Fisher’s exact test p-value = 0.42). We also looked at RNA expression relative to GTEx normal cervix tissue (n = 19) of proteins known to interact with PRDM9 as part of its binding to motifs and downstream proteins that it recruits^49,50^ (Fig. 6e). Although most of these proteins were not differentially expressed as compared to normal cervix tissue, we found that HELLS, an interacting protein of PRDM9 involved in its localization to motifs, exhibited significantly increased RNA expression in our cervical cancer cohort, as did RAD51, which is a marker of double-stranded break repair^50,51^. ZCWPW2 was downregulated in our cohort, but due to its redundancy with ZCWPW1 as a reader of PRDM9 marks^52^, we cannot be certain of the relevance of its downregulation. Our observations, alongside others’ work^48,49^ linking the motif to double-stranded breaks, raise the new possibility that PRDM9 may play a role in HPV-human hybrid ecDNA formation.

### Allele-specific ChIP-seq data can be used to investigate ecDNA enhancer and promoter landscapes

To improve understanding of ecDNA-specific epigenetic regulation, we sought to identify ChIP-seq reads aligning to ecDNAs, distinguishing these from ChIP Seq reads aligning to genomic loci. To achieve this, we focused on sequence variants detected in our long read data, seeking to identify those variants that were also in our short-read ChIP-seq data. We thus developed a novel approach to assign a subset of ChIP-seq reads, specifically those that overlap heterozygous SNVs, to alleles called in the long-read WGS data (Fig. 7a, see Methods). We then used these data to investigate enhancers and promoters in an allele sensitive manner, with respect to ecDNA loci.

**Figure 7.**
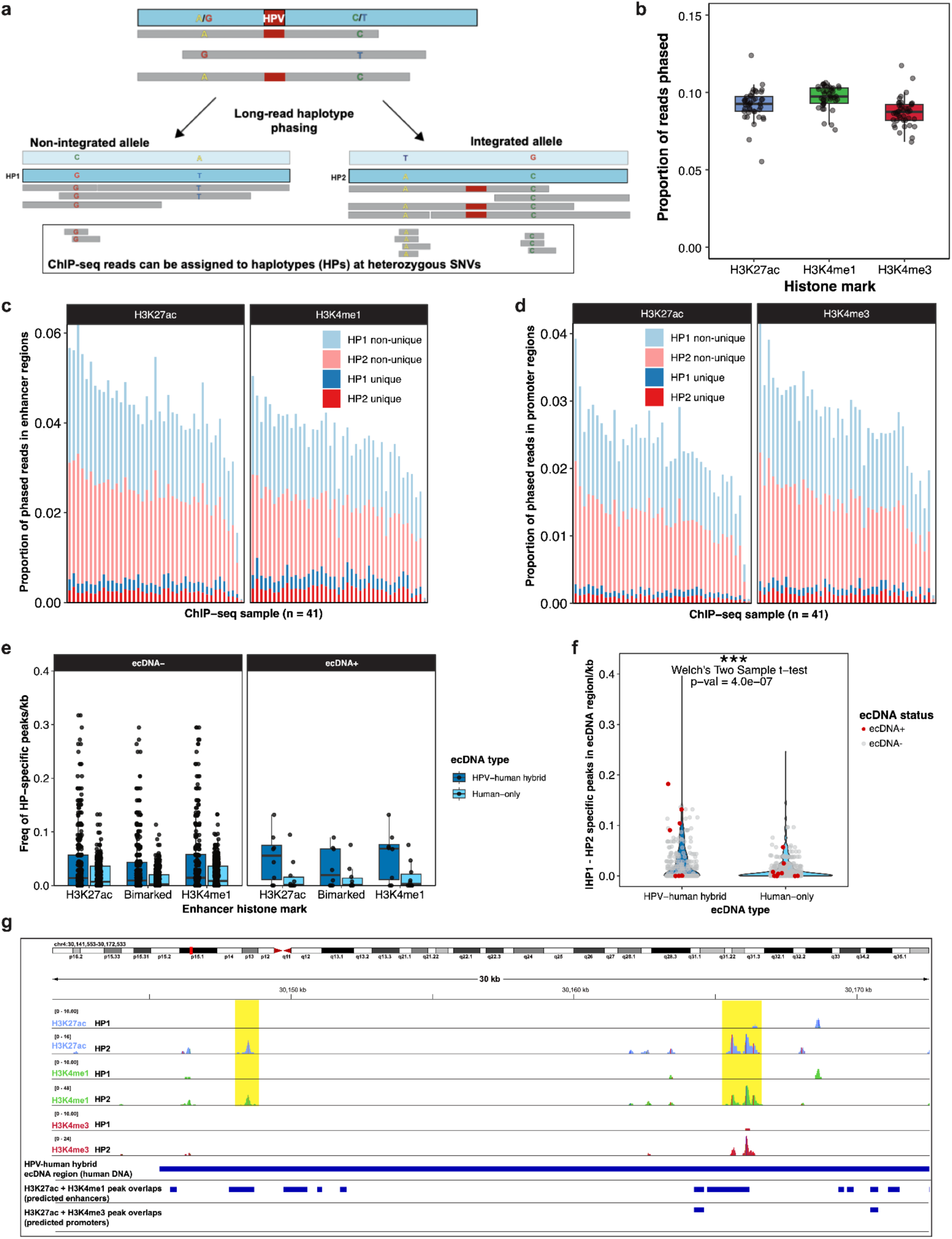
Using long-read WGS phasing to assign ChIP-seq reads to alleles reveals ecDNA-specific enhancers and promoters. **a** Simplified schema for assigning ChIP-seq reads to alleles at heterozygous SNVs. **b** Phased reads overlapping peak regions as a proportion of total unphased reads. Phasing metrics for ChIP-seq phasing pipeline (n = 41 samples, n = 3 marks for each sample) for proportion of reads overlapping enhancers (**c**) and promoters (**d**). **e** Frequency of allele-specific peaks per kb in ecDNA loci. **f** Allele (AL) bias in HPV-human hybrid ecDNAs compared to human-only ecDNAs. **g** Example of AL2-specific peaks in predicted enhancer loci within an HPV-human hybrid ecDNA locus.

Phased reads that fell in peak regions made up a median of 9.3% of total unphased reads (Fig. 7b). This corresponded to a median of 6.9 x 10^6^ reads (H3K27ac), 1.2 x 10^7^ reads (H3K4me1), and 5.7 x 10^6^ reads (H3K4me3) from 41 ChIP-seq bam files for each of three histone marks. We next overlaid these phased data with predicted enhancer or promoter peaks (defined as H3K27ac+H3K4me1 peaks or H3K27ac+H3K4me3 peaks respectively). We found that 3.2 x 10^6^ (4.3%) of H3K27ac reads and 4.6 x 10^6^ (3.4%) of H3K4me1 reads were able to be phased and overlapped predicted enhancer regions. Allele-specific H3K27ac or H3K4me1 peaks were found in 3.3 x 10^5^ (0.44%) and 4.1 x 10^5^ (0.58%) of enhancer regions respectively (Fig. 7c). 1.7 x 10^6^ (2.3%) of H3K27ac reads and 1.7 x 10^6^ (2.8%) of H3K4me3 reads were able to be phased and overlapped predicted promoter regions. There were 1.3 x 10^5^ (0.18 %) H3K27ac reads and 1.6 x 10^5^ (0.24%) of H3K4me3 reads in promoter regions that were allele-specific (Fig. 7d).

Summarizing allele-specific peaks in ecDNA regions revealed that allele-specific peaks in promoter or enhancer regions were not uncommon for ecDNAs. However, due to sample size constraints, it was difficult to determine if ecDNAs were enriched in allele-specific promoter or enhancer peaks compared to matched regions in ecDNA-samples (enhancers pictured in Fig. 7e). We expected ecDNA size and the number of allele-specific peaks to be correlated and indeed there were strong correlations observed for human-only ecDNAs (e.g. R between 0.85 to 0.91 and Pearson correlation p-value ≤ 0.002 for enhancer-specific peaks). However, there was no correlation between ecDNA size and the number of allele-specific peaks for HPV-human hybrid ecDNAs (e.g. R between 0.16 and 0.25 and Pearson correlation p-value > 0.05 for enhancer-specific peaks). To gain insight into whether allele-specific peaks in ecDNA regions were more commonly on one allele versus the other, which served as a proxy for potential ecDNA allele-specific peaks, we compared the absolute value of the difference in allele specific peaks in ecDNA regions controlled for ecDNA size. Allele bias was more pronounced for HPV-human hybrid ecDNAs compared to human-only ecDNAs (Welch’s two-sample t-test p-value = 4.0 x 10^-7^; Fig. 7f). These results support the notion that allele-specific peaks in enhancers and promoters on HPV-human hybrid ecDNAs are under selection. An example of ecDNA allele-specific peaks in enhancers is pictured in Fig. 7g.

## Discussion

We sought to characterize the structural and epigenetic organization of ecDNAs in cervical cancer to better understand the potential roles of these circular DNAs. Our study (1) characterized ecDNAs in Ugandan women with cervical cancer by integrating genomic, transcriptomic, and epigenomic profiling, (2) compared and contrasted different types of ecDNAs based on the presence or absence of HPV DNA, (3) profiled allele-resolved methylation patterns within ecDNA loci using long-read WGS data, and (4) isolated ChIP-seq reads in an allele-resolved manner to explore ecDNA-specific promoters and enhancers.

We found evidence that HPV-human hybrid ecDNAs and human-only ecDNAs are distinct, with notable structural and regulatory differences beyond simply the presence of HPV DNA. We predict that HPV-human hybrid ecDNAs are serving as promoter, enhancer and *E6*/*E7* vehicles, whereas human-only ecDNAs are serving as human oncogene vehicles. We found limited evidence in support of a publication that found ecDNA-specific oncogene promoter hypomethylation^22^. Overall, our study is important to understanding how both ecDNAs and HPV drive cervical cancer at an allele-specific level.

We hypothesize that HPV-human hybrid ecDNAs may not be serving as vehicles for human genes, but may instead be shuttling the HPV oncogenes, *E6* and *E7* and viral enhancers, as well as co-opting *trans*-activating human enhancers and promoters to drive viral gene expression. Consistent with this hypothesis, our study found that HPV-human hybrid ecDNAs contained a higher density of human promoters and viral enhancers relative to the corresponding loci in non-ecDNA containing samples and that *E6* expression was increased in HPV-human hybrid ecDNA containing samples compared to other samples in the cohort. Previous literature has described HPV-human hybrid ecDNAs as super-enhancer vehicles^23^, and our results do not refute this. We also found evidence that allele-specific promoter and enhancer peaks on HPV-human hybrid ecDNAs may be under selection, whereas the same cannot be said for human-only ecDNAs, but more research is needed to confirm this finding.

We also expanded on our earlier finding of allele-specific hypomethylation downstream of HPV integration on HPV-human hybrid ecDNAs^3^. Specifically, we found that these allelic hypomethylation regions were associated with H3K4me3 and H3K27ac peaks indicative of a putative human promoter region. All HPV-human hybrid ecDNAs with long-read WGS data, regardless of allelic hypomethylation status, had evidence for HPV-human fusion transcript expression. We note that these regions of allelic hypomethylation were specific to HPV-human hybrid ecDNA loci and were methylated on both alleles in comparison samples. Additionally, human genes falling within allelic hypomethylation regions were under ASE, thus confirming that epigenetic differences between alleles correlate with the transcriptome. Although we cannot determine the causal relationship between HPV hypomethylation and HPV-human hybrid ecDNA formation, this work supports recent analysis associating viral SV with changes to local epigenome profiles in cancer^58^.

Our study has a few technical limitations. We detected ecDNAs across the cohort using AmpliconArchitect, which relies on short-read WGS data. Notably, ecDNAs are large (∼100 kb-5 Mb) and therefore difficult to resolve unambiguously using short-read WGS data. Although we validated the majority of the AA results using long-read WGS data (*de novo* assembly with Flye and supplemental read support), we cannot be confident of the structures of the 15% of ecDNAs that failed to validate. We note that long-read ecDNA detection tools are emerging, which may improve upon ecDNA predictions in the future^59,60^. Another limitation of our study is the correlative nature of some of the ChIP-seq results and the possible confounding of high CN of ecDNAs. We address the former with phased ChIP-seq data derived from the long-read phased heterozygous SNVs and the latter with a CN adjustment step, but we also note that there may be biases in these data. For example, in the case of our phased ChIP-seq data, we cannot prove that the absence of reads from the remaining allele actually corresponds to the absence of signal and we note we may instead be limited by coverage of these specific loci.

In summary, our study compared and contrasted HPV-human hybrid and human-only ecDNAs in terms of structure and regulation in cervical cancer. We found that ecDNAs have allele-specific regulatory patterns, which is consistent with the notion that there exists differential regulation between ecDNAs and the underlying chromosomal DNA from which they arise, and provide evidence that the presence of HPV DNA within ecDNAs fundamentally changes the predicted function of such ecDNAs.

## Acknowledgements

We gratefully acknowledge the Fred Hutchinson Cancer Research Center for providing the HTMCP cervical cancer samples for genomic analyses. S.M. and V.L.P. were the recipients of CIHR Frederick Banting and Charles Best Canada Graduate Scholarships FBD-187583 and GSD-152374 respectively. M.N. was supported by postdoctoral fellowship awards from the Canadian Institutes of Health Research (FRN-188098) and Michael Smith Health Research BC (RT-2023-3168). This work was supported in part by funding provided by the Canadian Institutes for Health Research (CIHR award FDN-143288 and PJT-180410) to M.A.M. M.A.M is the Terry Fox Leader in Cancer Genome Science. This study was conducted with the financial support of The Terry Fox Research Institute and the Terry Fox Foundation. The views expressed in the publication are the views of the authors and do not necessarily reflect those of the Terry Fox Research Institute or the Terry Fox Foundation.

## Author contributions

This study was conceived by S.M. and M.A.M. Data were generated at Canada’s Michael Smith Genome Sciences Centre and analyses were performed by S.M., V.L.P., and R.D.C. M.A.M. supervised this work. S.M., M.N., P.P., Y.Z., and D.T. performed the ecDNA validation. S.M. wrote the original draft. All authors read and edited the manuscript.

## Data availability

The raw long-read whole-genome sequencing data have been deposited to dbGaP under the dbGaP study ID (phs003780.v1.p1). The long-read cDNA sequencing data is in the process of being deposited under the same dbGaP study ID. Other data from the HTMCP dataset used in this study have been deposited in dbGaP under the dbGaP study ID (phs000528) as part of the NCI Cancer Genome Characterization Initiative (CGCI; 613 phs000235). The processed expression data used in this study can be downloaded from the Genomic Data Commons (GDC) data portal (https://portal.gdc.cancer.gov/).

## Code availability

The figure generation code is available on GitHub: https://github.com/MarraLab/ecDNAs_cervical. Our bioinformatic tool: Separation of ChIP-Seq reads from phased long-read Mapped Alignments (SChISMA), which first phases long-read WGS data and then matches SNVs from ChIP-seq data to haplotypes called in the resultant long-read WGS vcf file, is available on GitHub: https://github.com/sigmaclennan/SChISMA.

## Source data

Available upon request.

## Supplemental Figures

**Supplemental Figure 1.**
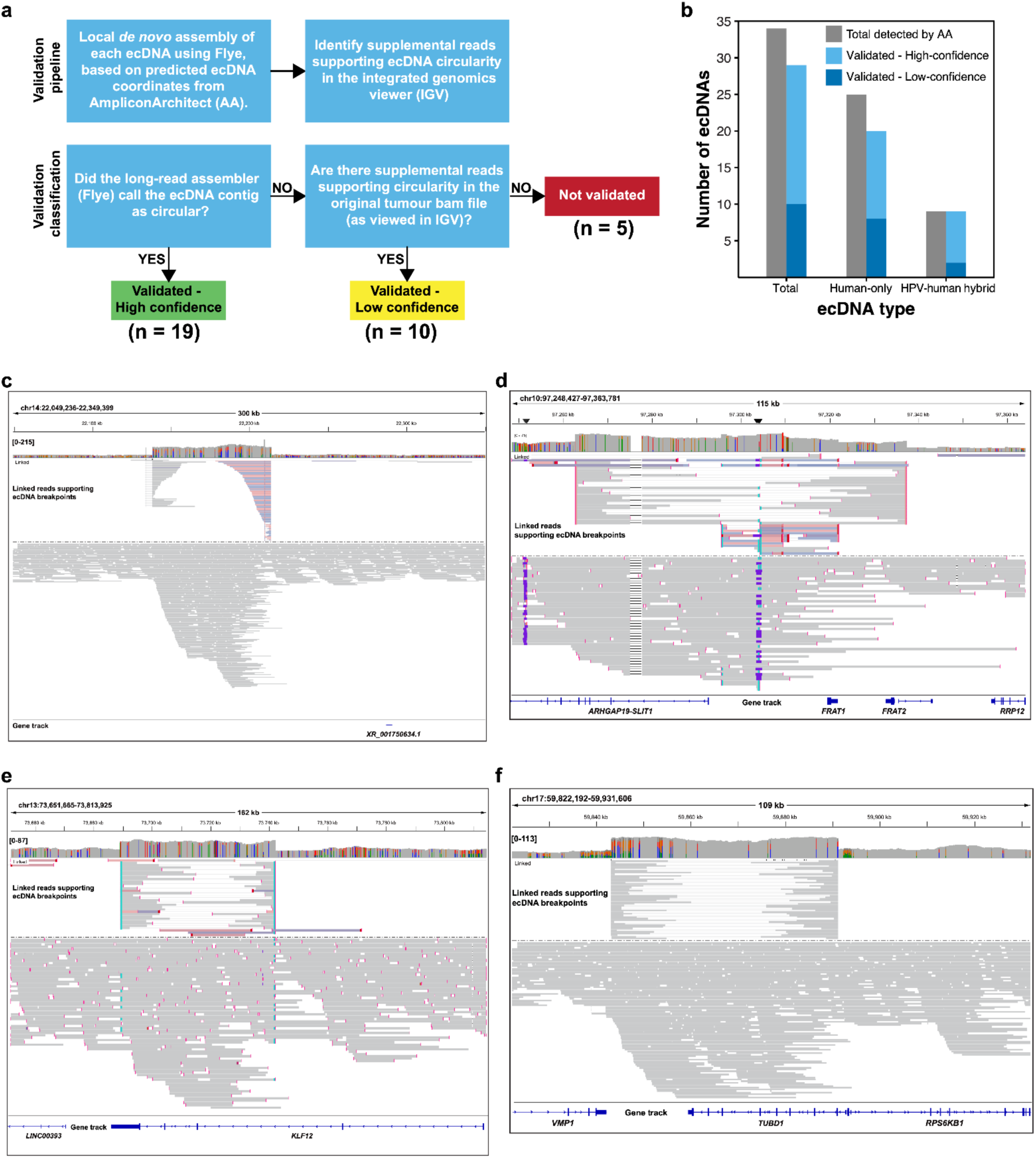
Validation of ecDNA circularity using a long-read WGS approach. **a** Long-read validation pipeline and classification scheme. **b** Long-read validation results. **c-f** Examples of low confidence ecDNAs predicted by AA and validated with supplemental read support in IGV (linked reads supporting ecDNA breakpoints are displayed at the top of each read picture).

**Supplemental Figure 2.**
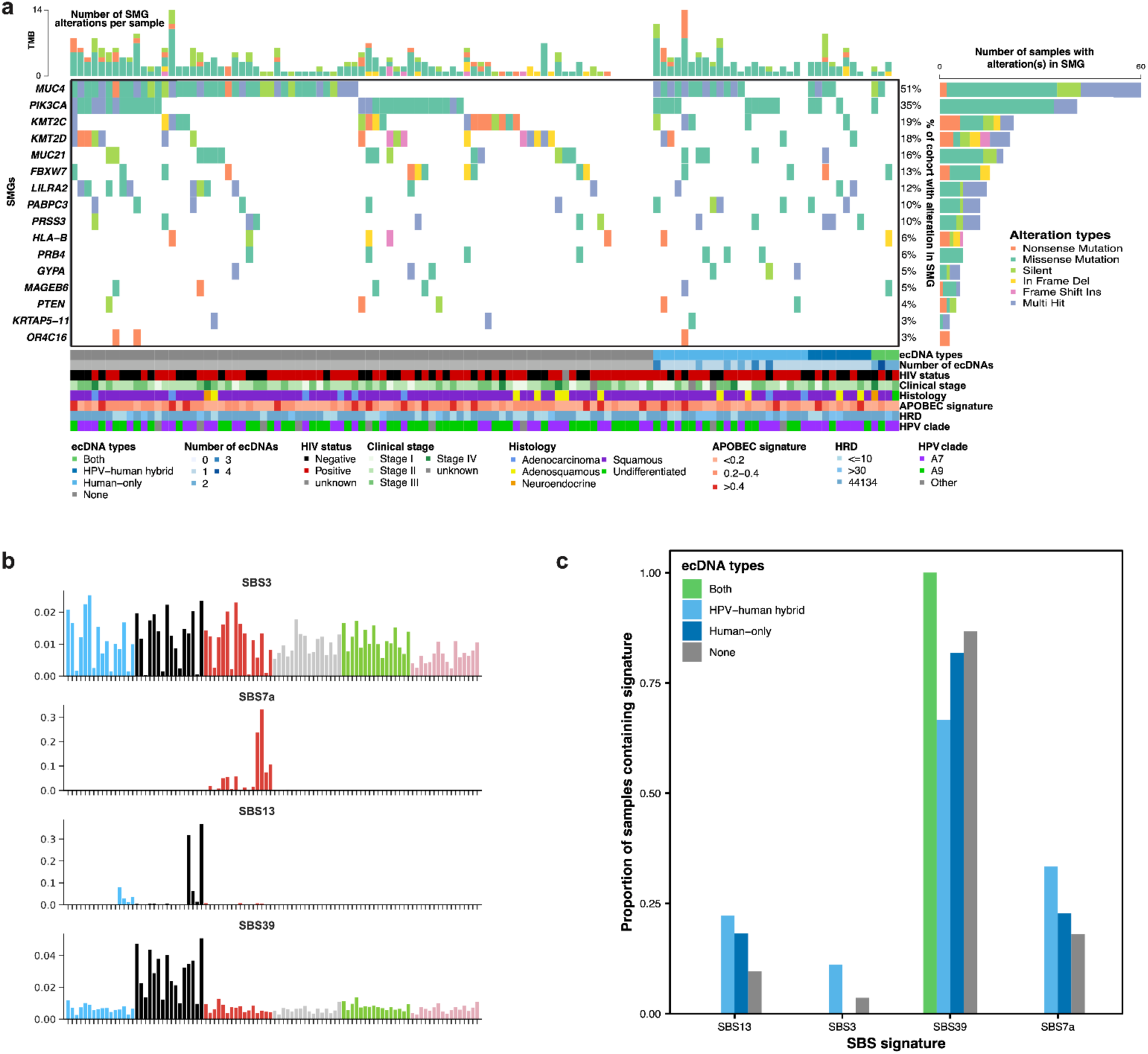
Correlation of cohort characteristics and molecular features with ecDNA presence and class. **a** Oncoplot showing ecDNA presence and other cohort characteristics and significantly mutated genes (SMGs). Oncoplot is ordered by ecDNA type. **b** Matched COSMIC SBS signatures from MuSiCal with SNVs called by Strelka2. **c** Proportion of samples of different ecDNA classes with the four matched signatures.

**Supplemental Figure 3.**
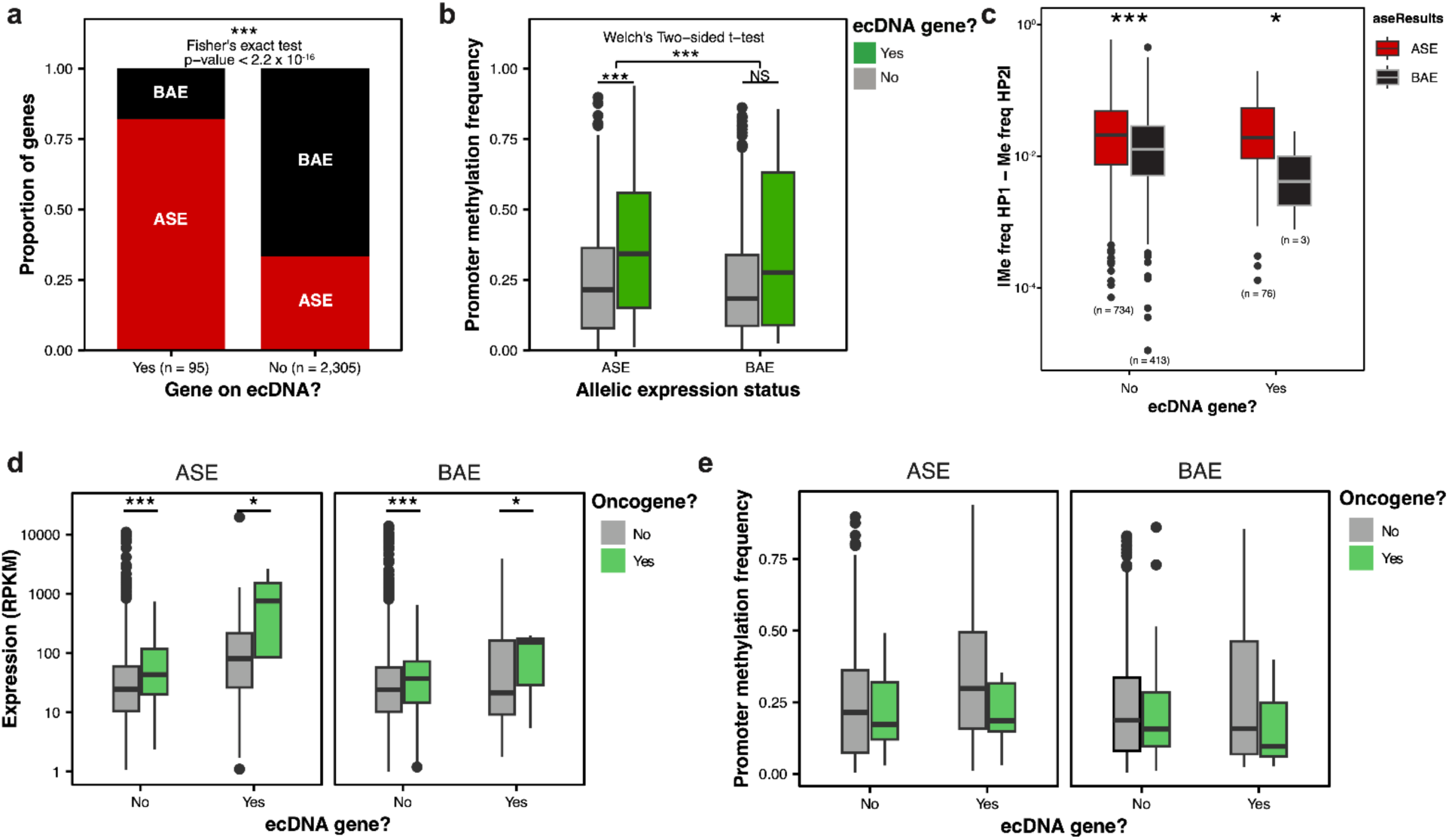
ecDNAs tend to bear genes that are allelically expressed and this is likely not driven by promoter hypomethylation. **a** Proportion of genes that display allele specific expression (ASE) or bi-allelic expression (BAE) comparing genes predicted to reside on ecDNAs to those not on ecDNAs for n = 46 samples with long-read WGS data available. **b** Promoter methylation frequency for ASE and BAE genes comparing genes on ecDNAs to those not on ecDNAs (n = 46 samples, n = 2,400 genes). **c** Methylation frequency (Me Freq) difference between alleles (ALs) for ecDNA genes and their allelic expression status. **d** Expression of oncogenes and non-oncogenes on ecDNAs relative to their allelic expression status. **e** Promoter methylation frequencies for oncogenes and non-oncogenes on ecDNAs relative to their allelic expression status.

**Supplemental Figure 4.**
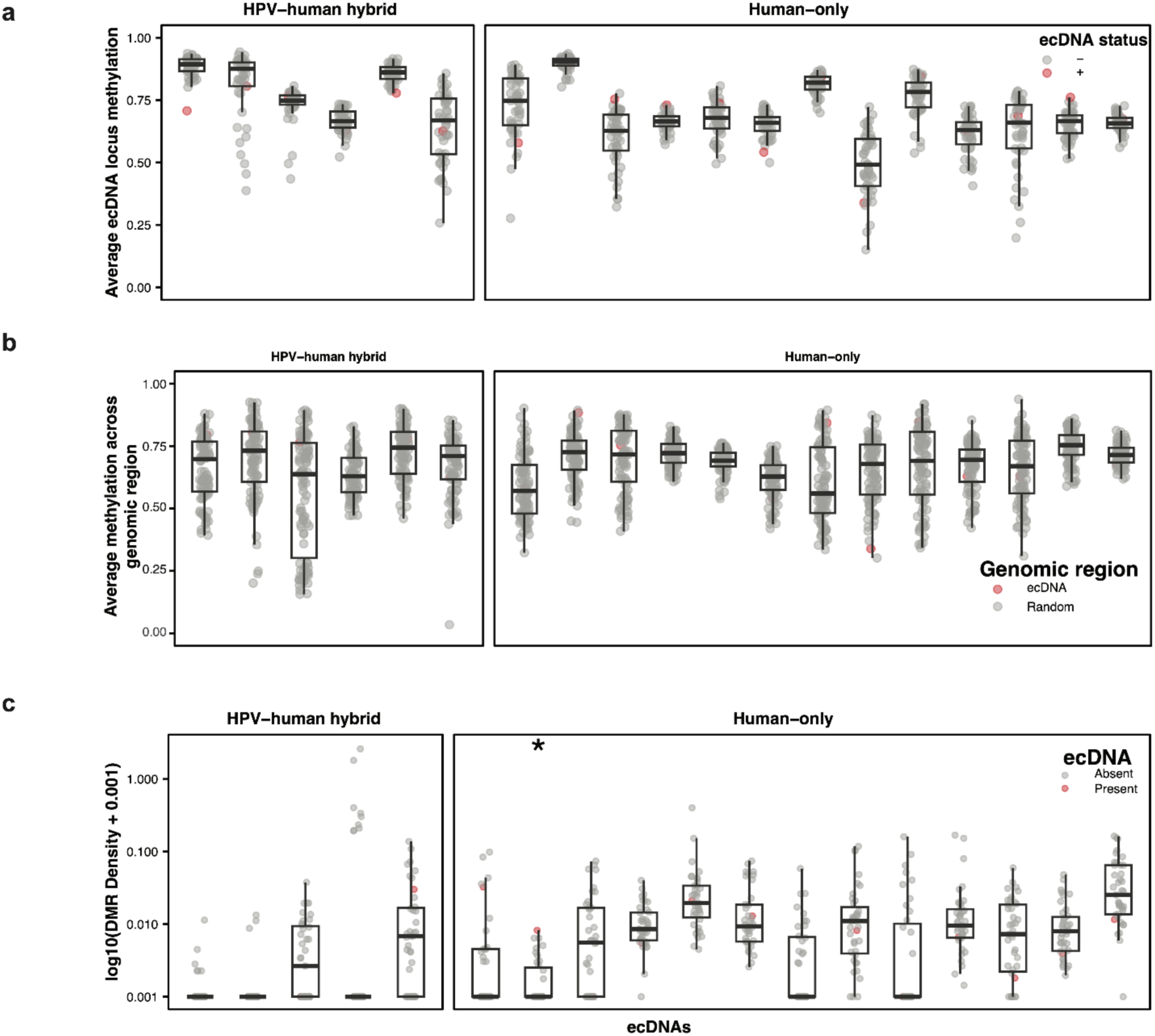
ecDNAs do not portray global methylation differences and are not enriched in allelic differentially methylated regions (aDMRs). **a** Average ecDNA locus methylation comparing the sample bearing the ecDNA (n = 1) to the rest of the long-read WGS cohort (n = 44), ANOVA permutation test with FDR < 0.05 multiple testing correction. **b** Average locus methylation comparing ecDNA loci (n = 19) to a random sample of genomic regions (n = 100), ANOVA permutation test with FDR < 0.05 multiple testing correction. **c** aDMR density within each ecDNA locus compared to the null distribution, * = adj. p-value < 0.05.

**Supplemental Figure 5.**
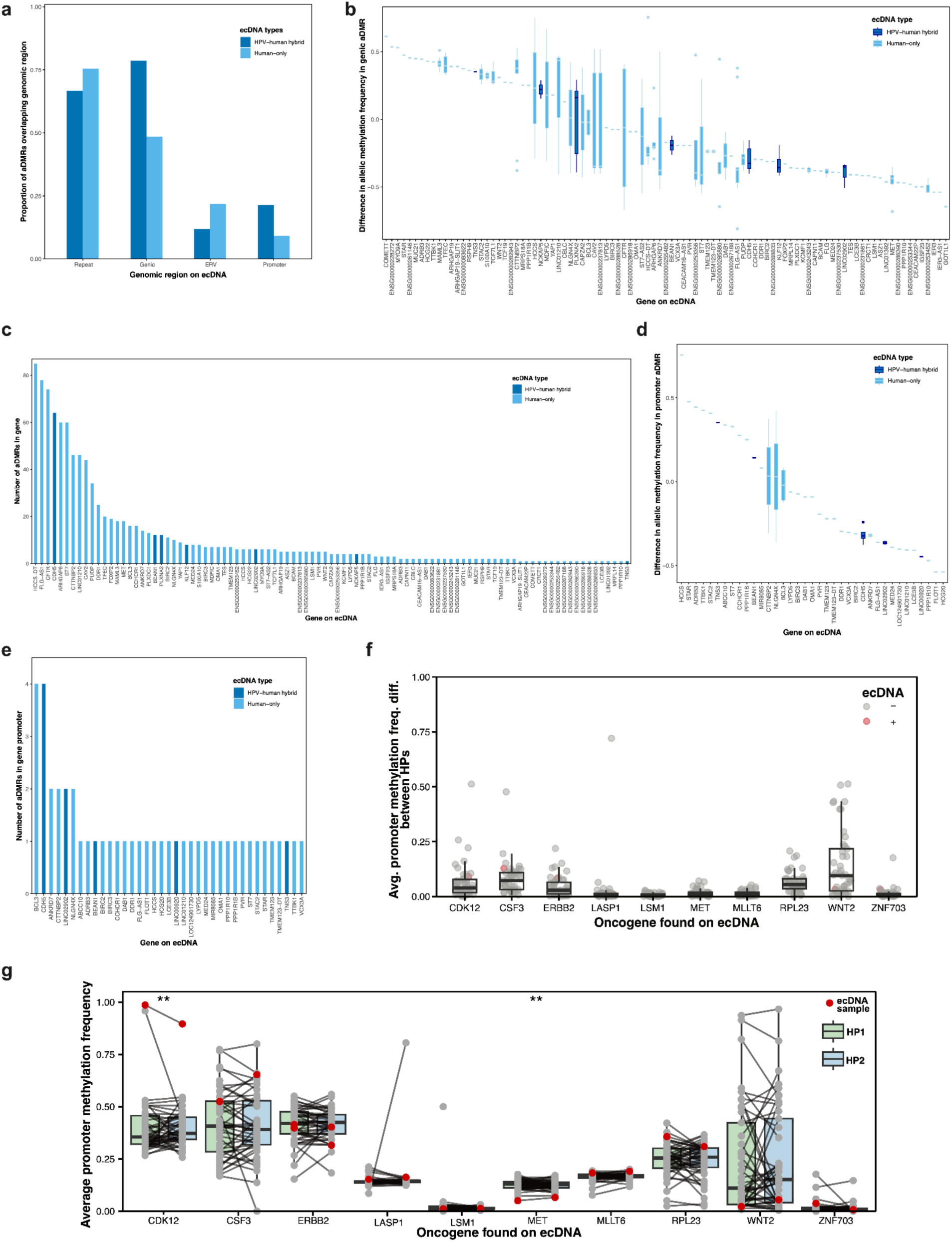
Analysis of ecDNA genic and promoter aDMRs. **a** Genomic regions covered by ecDNA aDMRs coloured by ecDNA type. **b** Difference in allelic methylation between alleles for genic aDMRs in ecDNA loci. **c** Number of aDMRs in genes in ecDNA loci by gene and ecDNA type. **d** Difference in allelic methylation between alleles for promoter ecDNA aDMRs. **e** Number of predicted ecDNA aDMRs in promoters by gene and ecDNA type. **f** Average promoter methylation frequency difference between alleles (ALs) for oncogenes found on ecDNAs (n = 47 samples). **g** Average promoter methylation frequency for each alleles in each sample with available long-read WGS data (n = 47) for oncogenes found on ecDNAs (ANOVA permutation test with FDR < 0.05 multiple testing correction; * = adj. p-value < 0.05, ** = adj. p-value < 0.01.

**Supplemental Figure 6.**
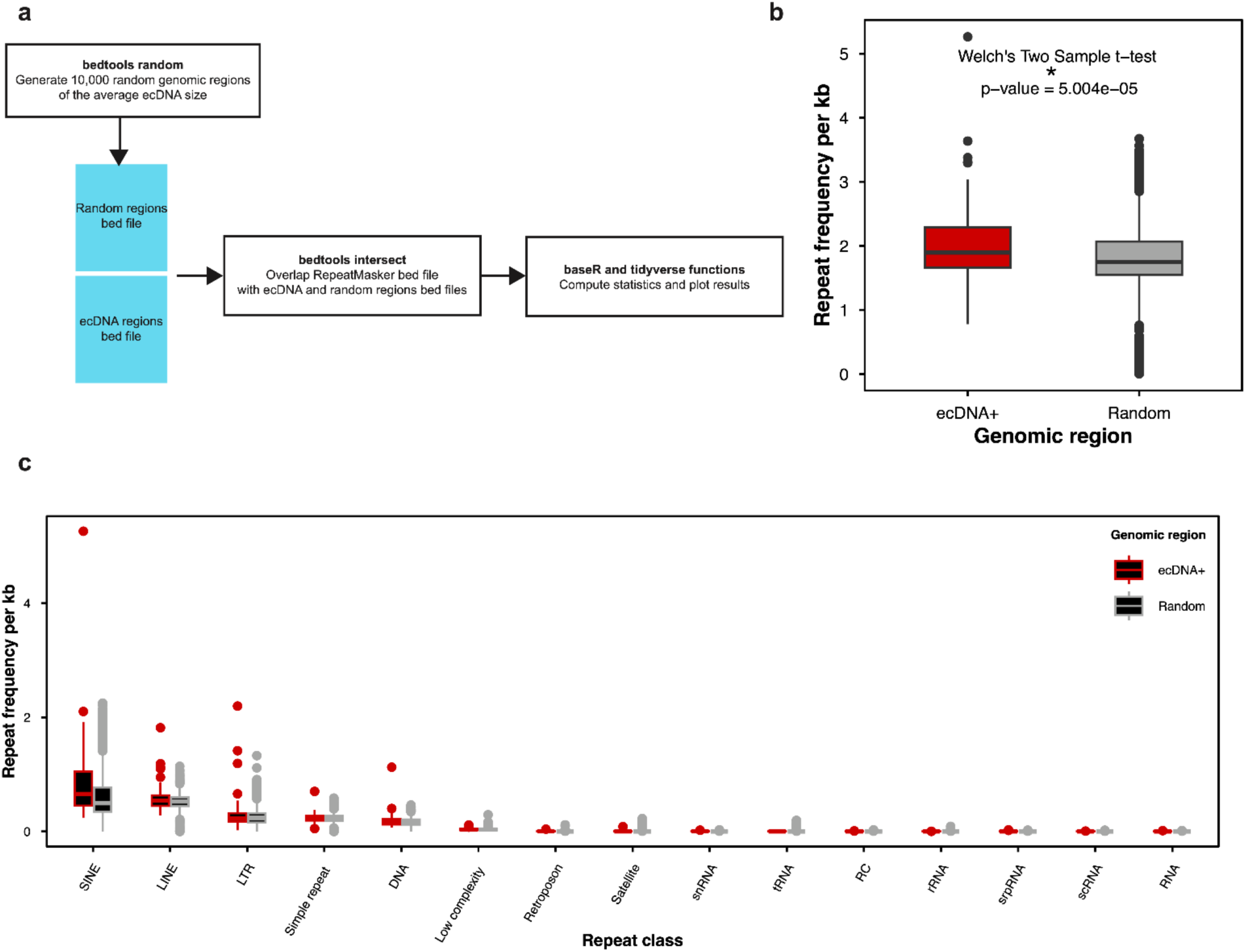
ecDNAs are enriched in repetitive regions compared to the rest of the genome, but not enriched in any particular repeat class. **a** Pipeline to compute and compare repeat frequencies in ecDNA loci to randomly generated genomic regions. **b** Comparison of repeat frequency on ecDNAs (n = 49) and a random sample of similarly sized genomic regions (n = 100, length = 181,254 bp). **c** Repeat classes represented on ecDNAs stratified by ecDNA type (n = 49). **c** Repeat classes represented on ecDNAs as compared to a random sample of similarly sized genomic regions (n = ∼10,000, length = 181,254 bp). * = p-value < 0.05.

**Supplemental Figure 7.**
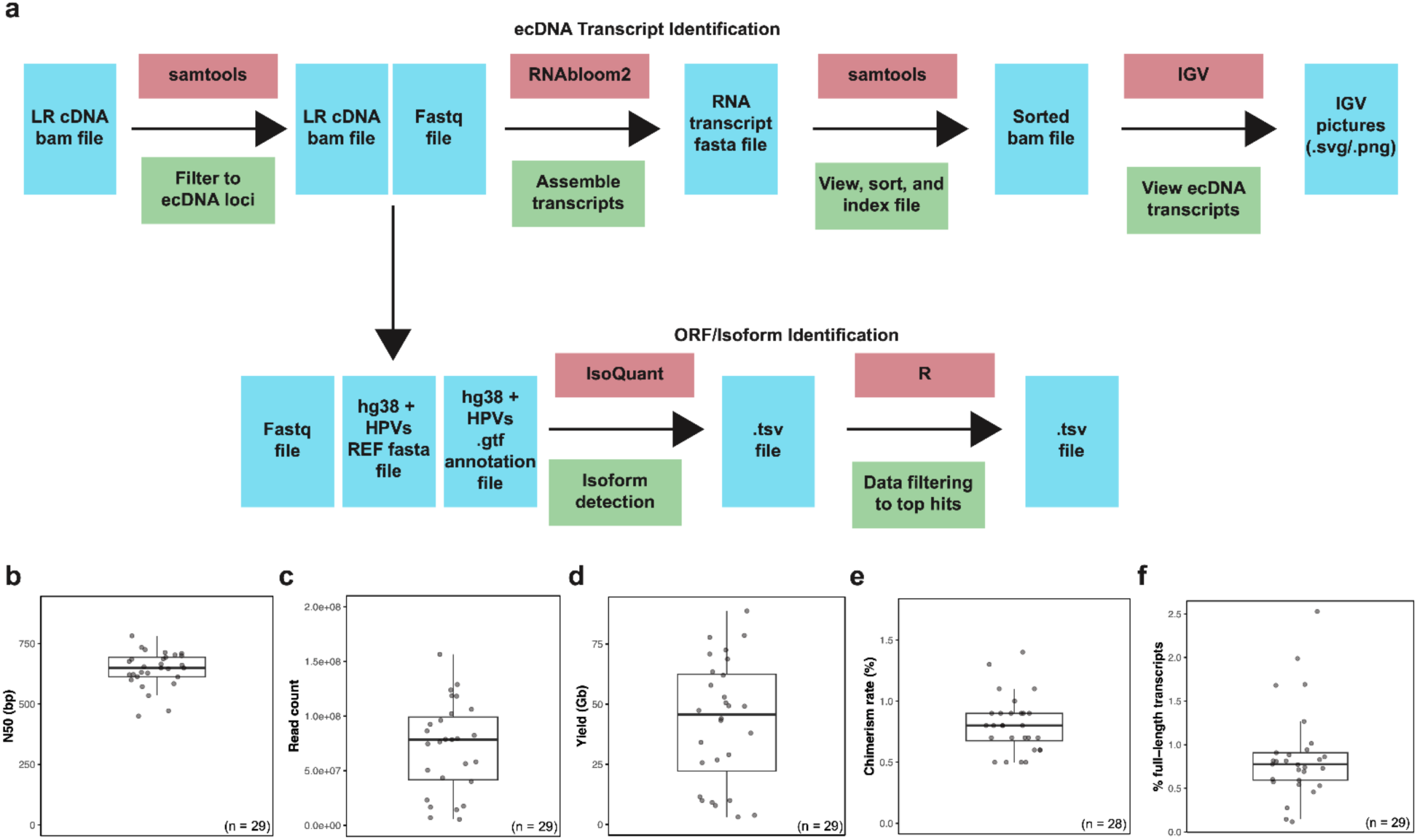
Workflow and quality control (QC) metrics for long-read cDNA-sequencing (LR cDNA-seq) dataset. **a** Workflow for long-read cDNA-seq analysis, QC metrics for long-read cDNA-seq dataset (n = 29): **b** N50, **c** Total read count, **d** Yield, **e** chimerism rate, **f** percentage of reads covering full-length transcripts.

## Supplemental Tables

Supp. Table 1 - High-confidence genes contained on ecDNAs as predicted by AmpliconArchitect

Supp. Table 2 - The allelic expression status of genes predicted to reside on ecDNAs in samples containing long-read WGS data

Supp. Table 3 - Genes and gene promoters overlapped by aDMRs in ecDNA loci

## Methods

### Ethical approval

Our study performs genomics analyses of samples approved for this use by the Fred Hutchinson Cancer Research Center Institutional Review Board (7662). Ethical approval was also obtained from the BC Cancer Research Ethics Boards (University of British Columbia (UBC) BC Cancer REB H16-02279) for molecular characterization as previously described^2^ and more recently by UBC (REB H24-02822).

### Pathology and molecular review

Formalin-fixed, paraffin-embedded (FFPE) tumour blocks were sent from the Uganda Cancer Institute to the University of California, San Francisco for histopathological review. Three pathologists reviewed the hematoxylin and eosin (H&E) and p16 immunohistochemistry (IHC) slides at the Nationwide Children’s Hospital as previously described^2^.

### Cervical cancer samples

Cervical cancer samples from the human immunodeficiency virus (HIV)+ Tumour Molecular Characterization Project (HTMCP) were profiled as part of this study^2^. Short-read paired-end whole-genome sequencing (WGS, Illumina HiSeq 250, n = 118^2^) and long-read WGS (Oxford Nanopore Technologies (ONT) PromethION, n = 47^3^) data, as well as whole-transcriptome sequencing (WTS/ RNA-sequencing (RNA-seq), Illumina HiSeq 2500, n = 118^2^) and native chromatin immunoprecipitation-sequencing (ChIP-seq) data (Illumina HiSeq 2500, n = 50^2^) were analysed in this study.

### Whole-genome sequencing

#### Library preparation

Short-read paired-end WGS library preparation for the cervical cancer samples (n = 118) is previously described^2^. Library preparation for the long-read WGS data involved size selection using the BluePippin method (SageSciences) and was previously published^3^.

#### Sequencing

Cervical cancer samples (n = 118) underwent short-read WGS with 125-bp paired-end reads to a target depth of 80X on the Illumina HiSeq 2500 as part of our previously published study^2^. The ONT PromethION sequencer was used to obtain long-read WGS data and methylation data on the cervical cancer samples (n = 72 for long-read WGS validation of which n = 47 were used for the methylation analyses, the complete protocol is described^3^).

#### Short-read RNA-sequencing / Whole-transcriptome sequencing

RNA-seq library preparation involved Poly(A) mRNA selection and cDNA synthesis of the cervical cancer samples (n = 118) and is described in detail in our previous study^2^. Short-read WTS (RNA-seq) was performed on the cervical cancer samples (n = 118) on the Illumina HiSeq 2500 as previously described with 75-bp paired-end reads^2^.

#### Native chromatin-immunoprecipitation sequencing

ChIP-seq library preparation for the cervical cancer samples is described in our previous study^2^. ChIP-seq libraries were normalized and pooled prior to sequencing on the Illumina HiSeq 2500^2^. ChIP-seq for six different histone marks (H3K4me1, H3K4me3, H3K27ac, H3K36me3, H3K9me3, and H3K27me3, plus an input control) was performed on 47 samples and an additional 5 samples had a subset of these six marks as previously described^2^. Fifty of these samples were analysed as part of this study.

#### Long-read cDNA sequencing

Long-read cDNA-seq library preparation for cervical cancer samples was using the method referenced^63^. Briefly, the method relies on rRNA-depletion and does not use gTube shearing to maintain transcript length. Sequencing following library preparation was done on an ONT PromethION sequencer.

#### Sequencing data pre-processing - alignment and basecalling

Cervical cancer WGS and WTS bam files were aligned to a HPV-human hybrid hg38 reference genome, using STAR RESM^64^ (v.2.5.2b) with an Ensembl100+HPV annotation (RNA-seq/WTS) and minimap2^65^ (v.2.15, short-and long-read WGS, ChIP-seq). Prior to alignment, Guppy 5^66^ (“super accurate model” setting) was used for long-read WGS base-calling.

#### ecDNA discovery in short-read WGS data

The extrachromosomal DNA (ecDNA) detection tool, AmpliconArchitect^24^ (AA, v.1.2), was used to identify putative ecDNAs in the cervical cancer samples (n = 118). Briefly, AA uses paired-end short-read WGS data as input and is a graph-based method for identifying ecDNAs as well as other focal amplification types^24^. The tool was run in a conda environment using CNVkit^67^ (v.0.9.10.dev0) for copy number calling with default settings using a hg38 reference genome with high-risk HPV types added as additional chromosomes. Results were visualized using a combination of R (v.4.0.2) packages (tidyverse^68^ (v.2.0.0)), and other tools (circos^69^ (v.0.69.9), IGV^27^ (v.2.17.0)).

#### ecDNA validation using long-read WGS

To validate the putative ecDNAs predicted by AA^24^ (v.1.2), we first utilized the long-read WGS data. We performed an additional validation of the 21 putative ecDNAs predicted by AA^24^ (v.1.2) with long-read WGS data available (n = 16 samples) using a combination of the *de novo* long-read assembly tool, Flye^26^ (v.2.9.2), and manual review of supplemental reads in IGV^27^ (v.2.17.0). Specifically, Flye^26^ (v.2.9.2) was run on fasta files cut according to ecDNA bedfile coordinates from AA^24^ (v.1.2) with an additional 5 kb of flanking regions surrounding each genomic region using samtools^70^ (v.1.9) commands (view, sort, and fasta) with three rounds of polishing from minimap2^65^ (v.2.24, --i 3) and the --meta mode. An ecDNA was considered high confidence if a circular contig was predicted by Flye^26^ (v.2.9.2) to reside within the coordinates of the bedfile output by AA^24^ (v.1.2), whereas an ecDNA was considered low confidence if Flye^26^ (v.2.9.2) did not predict a circular contig, but manual review in IGV^27^ (v.2.17.0) revealed read support for supplemental reads connecting the junctions of the ecDNA (i.e. consistent with circularity). We checked that HPV was contained in the circular contigs from AA predicted HPV-human hybrid ecDNAs using seqkit^71^ (v2.4.0) to filter to the contig of interest, minimap2^65^ (v.2.24; with “-ax asm5” commands) to map the contig to the human-HPV hybrid reference sequence, and the samtools^70^ (v1.9) commands: view, sort, and index to generate a contig bam file. Resulting bam files were inspected in IGV^27^ (v.2.17.0) to confirm the presence of HPV reads.

#### Parts of HPV contained on HPV-human hybrid ecDNAs

To obtain what parts of HPV were predicted to occur on HPV-human hybrid ecDNAs, we utilized the bedfile output files from AA^24^ (v.1.2). Additionally, for one sample with multiple predicted substructures, we looked at predicted discordant reads between human and HPV DNA as part of the “sample#_amplicon#_graph.txt” output file from AA that was consistent with the putative HPV-human hybrid substructure.

#### Oncogene expression from putative ecDNAs

The oncogenes on ecDNAs were identified from the AA^24^ (v.1.2) output, which is based on the catalogue of mutations in cancer (COSMIC) annotation^72^. Evidence of oncogene amplification for genes on ecDNAs was found using RNA-seq data. Briefly, we processed samples by computing reads per kilobase of transcript per million mapped reads (RPKM) using the limma^73^ (v.3.46.0) and tidyverse^68^ (v.2.0.0) R packages. Statistically, we used a permutation test in R to compare RPKM for the cervical cancer sample bearing the oncogene on a ecDNA to the rest of the cohort with false discovery rate (FDR) multiple testing correction < 0.05 using the coin^74^ (v.1.4-3) R package.

#### Gene set enrichment analysis (GSEA)

Oncogenes found on ecDNAs (n = 33) were run through GSEA using the enrichR^75^ (v.3.2) package and the KEGG 2016^76^ gene annotation to identify significant pathway enrichments. Results were plotted using tidyverse^68^ (v.2.0.0) functions.

#### Significantly mutated genes (SMGs)

Somatic single nucleotide variants (SNVs) and indels were identified using Strelka2^31^ (v.2.9.10) for the cohort as compared to matched normal blood samples (n = 118). Subsequently, sample mutation calls were combined into a single formatted dataframe and were run through dNdScv^29^ (v.0.0.1.0) using default settings to identify significantly mutated genes (SMGs). A comparison of SMGs in ecDNA+ and ecDNA-samples was also performed using the maftools^77^ (v.2.6.05) mafCompare function in R (v.4.0.2). An oncoprint showing SMGs stratified by ecDNA presence was constructed using maftools^77^ (v.2.6.05).

#### Allele specific expression (ASE) of ecDNA genes

To identify whether ecDNA genes presented with allele specific expression (ASE), we utilized the Integrated Mapping and Profiling of Allelically-expressed Loci with Annotations (IMPALA)^36^ tool. Specifically, we ran the tool using the phasing option (results obtained from WhatsHap^78^ (v.1.0)) on samples with RNA-seq and long-read WGS data available, excluding one sample (n = 46). Next, we filtered the gene list for each sample to only the genes found to be present on ecDNAs in these samples (n = 95) and compared ecDNA genes to these same genes in samples not predicted to contain ecDNAs with these genes (n = 2,305). To compare promoter methylation values, we used the annotatr^79^ (v.1.16.0) “hg38_basic_genes” annotation filtered to promoter regions and extracted average CpG methylation results for each of these genes from Nanopolish^80^ (v.0.13.2) results. Annotations from oncogenes and non-oncogenes were obtained from the AA^24^ output (which is based on a COSMIC^72^ annotation). For allele specific methylation comparisons between ASE and BAE genes (ecDNA+/-) we utilized phased methylation data from NanoMethPhase^81^ (v.1.0) and then calculated the average methylation frequency across promoters annotated with annotatr^79^ (v.1.16.0) for each allele (AL), as described above. Results were plotted using tidyverse^68^ (v.2.0.0) functions.

### ecDNA methylation

#### Long-read WGS preprocessing - Structural variant calling, methylation calling, and phasing

Structural variants (SVs) were called using Sniffles^82^ (v.1.0.12b). Clair3^83^ (v.0.1-r8) was used to call small variants, which were subsequently phased into alleles using WhatsHap^78^ (v.1.0). Phase blocks were saved for subsequent analysis. Nanopolish^80^ (v.0.13.2) was used for methylation calling followed by NanoMethPhase^81^ (v.1.0) to phase methylation calls into alleles.

#### Global ecDNA methylation

Global ecDNA methylation values (not allelically resolved) across ecDNA loci was determined by computing the mean methylation frequency across all CpG sites within each ecDNA locus. Specifically, ecDNA bedfiles from AA^24^ (v.1.2) were used to filter methylation frequency call files from Nanopolish^80^ (v.0.13.2) to ecDNA coordinates. These results were compared to the same loci in non-ecDNA containing samples (n = 46) as well as randomly generated genomic regions. Specifically, the bedtools^84^ (v.2.27.1) “random” command was used to generate 100 genomic regions the same size as each ecDNA (n = 19). Methylation frequencies were computed for each of the 100 genomic regions and those lacking CpG sites were excluded (sample sizes varied slightly between ecDNAs). Plots for both comparisons were generated using tidyverse^68^ (v.2.0.0) functions following permutation tests using the coin^74^ (v.1.4-3) R package with multiple testing correction as FDR < 0.05.

#### Allelic differentially methylated regions (aDMRs)

To identify allelic differentially methylated regions (aDMRs) between tumour alleles, the DSS^85^ (v2.38.0) R package was used. Subsequently, a custom python script was used to determine enrichment of aDMRs within ecDNA loci as compared to a null distribution^3^ (https://github.com/vanessa-porter/Porter-ONT-HPV-FigureScripts2023). To determine the genomic regions covered by aDMRs, the bedtools^84^ (v.2.27.1) “intersect” function was used for annotations for genes (GENCODE (v.42)^86^, downloaded from UCSC Table Browser^87^ on Jan. 30, 2023), endogenous retroviruses (ERVs, downloaded from: https://herv.img.cas.cz/ on Mar. 27, 2024^88^), repeats (from RepeatMasker^89^, downloaded from UCSC Table Browser^87^ on Jan. 27, 2023), and CpG islands (downloaded from UCSC Table Browser^87^ on Jan. 27, 2023). aDMRs were also queried against promoter regions from the annotatr^79^ (v.1.16.0) R package’s “hg38_basic_genes” annotation filtered to promoter regions by using the GenomicRanges^90^ (v1.42.0) R package.

#### Predicting ecDNA alleles

To predict ecDNA alleles, ecDNA-associated SVs were extracted from the AA^24^ (v.1.2) output (specifically the “sample#_amplicon#_SV_summary.tsv” files filtered to “ecDNA” features) and matched to the Sniffles^82^ (v.1.0.12b) SV calls to extract read names supporting these SVs. Read names were then mapped to NanoMethPhase^81^ (v.1.0) phased bam files using the samtools^70^ (v.1.9) “view” command combined with an awk command (see provided code). Mapped reads were visualized in IGV^27^ (v.2.17.0) with the allele containing mapped reads considered to be the predicted ecDNA allele. Further support for an ecDNA allele came from examining the coverage of all phased reads and seeing substantially higher read coverage in the predicted ecDNA allele. Phase blocks from WhatsHap^78^ (v.1.0) were also examined to see if phase block switching was likely to play a role within the ecDNA locus (i.e. the ecDNA assigned as different alleles in different regions). In Figure 4, we note that due to a phase block switch in the ecDNA region, the ecDNA allele was determined to be allele 2 (AL2) in the 5’ end and allele 1 (AL1) in the 3’ end (Fig. 4a). The two aDMRs fell within the *C7orf65* gene promoter, with the ecDNA allele (AL1) being highly hypo-methylated relative to the chromosomal DNA allele (AL2; Fig. 4b-c).

#### Oncogene promoter methylation in ecDNA loci

We compared average oncogene promoter methylation in ecDNA loci for samples predicted to contain an ecDNA bearing an oncogene to all other samples (n = 47) for the 10 whole oncogenes predicted to reside in ecDNA loci with long-read WGS data available. Specifically, we used the annotatr^79^ (v.1.16.0) “hg38_basic_genes” annotation filtered to the promoter regions of these 10 oncogenes. Plots were generated using tidyverse^68^ (v.2.0.0) functions following permutation tests using coin^74^ (v.1.4-3) with multiple testing correction as FDR < 0.05. Samples lacking methylation calls in oncogene promoter regions were excluded.

#### ecDNA type and class genomic comparison

Genomic features of ecDNA types (i.e. human-only vs HPV-human hybrid) were compared. Sizes of ecDNAs were extracted from the bedfile output files from AA^24^ (v.1.2). For the genes contained on ecDNAs, these were extracted from “sample#_amplicon#_gene_list.tsv” files for ecDNA amplicons filtering the “feature” column to “ecDNA”. Notably, genes that were not shared between multiple substructures were excluded from these lists, which is a limitation of this approach.

The number and types of SVs associated with each ecDNA were extracted from the “sample#_amplicon#_SV_summary.tsv” files again filtering the “feature” column to “ecDNA”. To determine the genomic region types covered by ecDNA loci, we used the bedtools^84^ (v.2.27.1) “intersect” function to query bedfiles output from AA against genes predicted by GENCODE^86^ (v.42; downloaded: Jan. 30, 2023), CpG islands (downloaded: Jan. 27, 2023), and repetitive regions from RepeatMasker^89^ (downloaded: Jan. 27, 2023) all downloaded from the University of California, Santa Cruz (UCSC) Table Browser^87^. Repeats overlapping ecDNA loci were also compared to the RepeatMasker^89^ overlap of a random sample of 10,000 genomic regions (with chromosomes other than 1-22, X removed) each the size of the median ecDNA in our cohort (181,254 bp) as generated by the bedtools^84^ (v.2.27.1) “random” command.

We compared the SNV mutational signatures of each ecDNA class (human-only, HPV-human hybrid, both, or none), by running MuSiCal^30^ (v.1.0.0) on genome-wide SNVs called with Strelka2^31^ (v.2.9.10) for each ecDNA class separately. We then computed the proportion of samples containing evidence for each of the significant signatures detected by MuSiCal.

We also compared ecDNA types in terms of whether the predicted ecDNA region overlapped a chromothriptic region. To call chromothripsis across the 72 samples with long-read WGS data, we used ShatterSeek^39^ (v.1.1) in R with CN calls from Ploidetect^91^ (v.1.4.2) and SV calls from Sniffles2^62^ (v.2.0.7). Next, we overlapped the significant chromothriptic regions (stratified by high and low confidence) with ecDNA regions separated by ecDNA type and made comparisons (Welch’s two-sided t-test with FDR < 0.05 multiple testing correction).

#### Motif analysis in HPV-human hybrid ecDNA loci

We also investigated motifs on HPV-human hybrid ecDNAs (n = 8) by looking at H3K4me1+H3K27ac (enhancers) and H3K4me3+H3K36me3 overlapping peak regions in HPV-human hybrid ecDNA regions using motifmatchr^92^ (v1.24.0). These peaks were originally called from MACS2^93^ (v.2.2.9.1), overlapped with a bedtools^84^ (v.2.27.1) “intersect” command, and cut to ecDNA regions prior to input into R (v.4.3.2). Focussing on the H3K4me3+H3K36me3 screens, we selected top hits by filtering to hits found in at least 35% of tested regions (30/85), and found to be significantly enriched as compared to nucleotide and size matched regions. We later focussed our analysis on the longest motif (MA1723.2; 20 bp) matching these criteria. We also inputted complete ecDNA loci (n = 13) as a test to see if motifs were enriched on ecDNAs more generally. We also overlaid these results with expression data from short-read RNA-seq as well as methylation data from long-read WGS.

### HPV-human fusion transcripts originating from ecDNAs

#### Quality control analysis

To assess the quality of our long-read cDNA-seq data, we calculated the chimerism rate and enumerated the yield, number of reads, and N50 for our dataset (n = 29, n = 28 for chimerism due to differing numbers of lanes). We also used BamSlam^94^ (commit 5d8ce25) to predict the number of full-length transcripts covered by single long reads. Specifically, we converted the aligned bams back to fastq files with the samtools^70^ (v.1.9) fastq command, randomly downsampled them, aligned them to the transcriptome (containing hg38 + high-risk HPVs) using minimap2^65^ (recommended settings for BamSlam - “minimap2-ax map-ont --sam-hit-only transcriptome.fasta reads.fastq | samtools view-bh > aligned_reads.bam”), and ran BamSlam^94^ with default settings. To quantify isoforms deriving from ecDNA loci, IsoQuant^95^ (v3.6.3) was run on fastq files cut to ecDNA regions (using samtools^70^ (v.1.9) commands) using the --data_type nanopore command and a gtf file for the hg38+HPV reference fasta file. Results were visualized in IGV^27^ (v.2.17.0) and further analyzed in R (v.4.3.2).

#### Detection of HPV-human fusion transcripts and other ecDNA transcripts

To detect HPV-human fusion transcripts from long-read cDNA-seq data we ran RNA-bloom2^96^ (v.2.0.1) in a conda environment on ecDNA regions (cut from bam files and converted back to fastq files using samtools^70^ (v.1.9) commands). Resulting transcript fasta files were aligned to the hg38+HPV reference fasta file using minimap2^65^ (output as sam file) and converted to a sorted bam file with samtools^70^ (v.1.9) commands. Results were visualized using IGV^27^ (v.2.17.0). Graphs were plotted using R (v.4.3.2) and tidyverse^68^ (v.2.0.0) functions.

### ChIP-seq analyses

#### De novo discovery of enhancers and promoters

To discover predicted enhancers and promoters in ecDNA loci de novo, we first called peaks with MACS2 (v.2.2.9.1) for H3K27ac, H3K4me1, and H3K4me3. MACS2 was run in a conda environment against each sample’s input control for each of the three marks for each sample containing ChIP-seq data (n = 50) with the-g hs,-f BAM, --broad (excluding H3K4me3), --broad-cutoff 0.1 (excluding H3K4me3), --no-model, and --extsize 147, and-q 0.01 (H3K4me3 only) commands. Peaks were further filtered to exclude contigs outside of chromosomes 1-22, and X and ENCODE blacklist regions for hg38^97^ (hg38-blacklist.v2.bed.gz, downloaded Feb. 6, 2026 from https://github.com/Boyle-Lab/Blacklist/tree/master/lists). Remaining peaks were filtered further to adjusted q-value < 0.01 (H3K27ac, H3K4me1) or adj. q-value < 0.001 (H3K4me3), fold enrichment of ≥ 3, 4, or 5 for H3K4me1, H3K4me3, and H3K27ac respectively, size of the peak region (150-1,500 bp for H3K4me3, 500-10,000 bp for H3K27ac, and 500-20,000 bp for H3K4me1). Peaks were also adjusted to account for local CN effects, by dividing signal values with absolute CN estimates from Ploidetect^91^ (v.1.4.2). Enhancers and promoters were predicted from overlaying H3K27ac and H3K4me1 and H3K27ac and H3K4me3 peaks respectively using the bedtools “intersect” function for unique overlaps. We then cut the resulting bed files to match the ecDNA loci according to the bedfile output from AA and compiled each ecDNA into a list of peaks from the sample predicted to contain an ecDNA in that region to the rest of the samples in order to run additional analyses in R (v.4.3.2) using tidyverse (v.2.0.0) functions. The density of enhancers or promoters per ecDNA was computed for all ecDNA+ samples compared to matched regions in ecDNA-samples.

#### Annotated enhancers

Evidence of annotated enhancer activity residing within ecDNA loci was determined using the coverage of H3K27ac as profiled using ChIP-seq (n = 50) for annotated enhancers from Genehancer^43^ (Dec. 2013 hg38 release, downloaded from UCSC Table Browser^87^: Mar. 11, 2024). To determine which annotated enhancers overlapped ecDNA loci, bedtools^84^ (v.2.27.1) “intersect” function was used querying ecDNA bedfiles from AA^24^ (v.1.2) with Genehancer^43^ (Dec. 2013 hg38 release) enhancers. Bigwig and bedgraph files were generated using the “bamcoverage” function from deepTools^98^ (v.3.5.2) using default parameters and the “--normalizeusing CPM” parameter. Results were visualized in IGV^27^ (v.2.17.0, bigwig files) and R (v.4.0.2, bedgraph files). Average/mean enhancer activity across ecDNA loci vs these same loci was calculated and statistically compared in R (v.4.0.2) using permutation tests with false discovery rate (FDR) multiple testing correction < 0.05 using the coin^74^ (v.1.4-3) R package for both HPV-human hybrid and human-only ecDNAs. Results were plotted using tidyverse^68^ (v.2.0.0) in R (v.4.0.2) and circos^69^ (v.0.69.9).

#### Phased ChIP-seq analysis

To better isolate ChIP-seq signals as deriving from alleles from which ecDNAs arise, we developed a novel technique to assign ChIP-seq reads to alleles by using long-read allele phased variants as a scaffold. The basic pipeline involves first phasing a given sample’s long-read WGS bam file using WhatsHap^78^ (v2.2) and also calling SNVs with Strelka2^31^ (v2.9.2) in the --exome mode (for expected uneven coverage) for a given histone mark in a ChIP-seq bam file from the same sample that is aligned to the same reference genome (in this case hg38 with high-risk HPV types added as additional chromosomes, although HPV is not “phase-able” with this technique).

Next, pysam (v0.23.3; https://github.com/pysam-developers/pysam) is used to extract read names from the resultant ChIP-seq SNV VCF for both REF and ALT alleles. Subsequently, ChIP-seq REF and ALT with read name information and long-read WGS phased VCF files are loaded into R for filtering. Specifically, variants from the ChIP-seq files that overlap phased variants in the long-read WGS VCF file are saved with their allele information (from long-read WGS) and their read names information (from ChIP-seq). Only read names are extracted from the resulting phased variants called in the ChIP-seq data and unique read names are saved in allele 1 (AL1) and AL2 files, which are subsequently used to filter the larger original ChIP-seq bam file into two allele-specific bam files using the samtools view-N [readnames]-b [output bam file] command. Resultant allele bam files are then sorted and indexed with samtools for viewing in IGV^27^ (v.2.17.0). We also analyzed HP-specific enhancers and promoters using these allele ChIP-seq bam files by using bedtools intersect commands for allele ChIP-seq reads overlapping H3K27ac and H3K4me1 (enhancers) or H3K4me3 (promoters) peaks called in the original ChIP-seq bam file (as part of the *de novo* discovery of enhancers and promoters analysis), as well as overlapping annotated enhancer loci from Genehancer^43^ (Dec. 2013 hg38 release, downloaded from UCSC Table Browser^87^: Mar. 11, 2024) with bedtools intersect or promoter loci from annotatr^79^ (v.1.16.0) “hg38_basic_genes” annotation filtered to promoter regions in R (v.4.3.2).

#### Promoter activity flanking HPV integrants in ecDNA loci

Coverage flanking (+/-10 kb) HPV for samples containing putative HPV-human hybrid ecDNAs and ChIP-seq data (n = 8) were extracted from bedgraph ChIP-seq files for six profiled histone marks (H3K4me1, H3K4me3, H3K9me3, H3K27ac, H3K27me3, H3K36me3) as generated by the “bamcoverage” function from deepTools^98^ (v.3.5.2) using default parameters and the “--normalizeusing CPM” parameter. ChIP-seq bedgraph ranges were converted into bp level signals using an awk command (see provided code). High coverage of H3K27ac and H3K4me3 (relative to H3K4me1) was taken as evidence for a promoter region.

